# Localized *Leishmania major* infection causes systemic iron deficiency that can be controlled by oral iron supplementation

**DOI:** 10.1101/2022.06.03.494645

**Authors:** Sourav Banerjee, Rupak Datta

**Author notes:** To whom correspondence to be addressed. Rupak Datta; Extn: 1214. Department of Biochemistry, University of Cambridge, United Kingdom (Sourav Banerjee).

## Abstract

*Leishmania major (L. major)* and its related parasitic species infection causes human cutaneous leishmaniasis that results into disfiguring skin lesion. Although *L. major* infection has been found to alter macrophage iron homeostasis we have limited understanding on whether it can also manipulate the same at systemic level. In fact, localized *L. major* infection found to promote iron deficiency anemia in children by an unknown mechanism. To address these unresolved issues, Balb/c mouse were infected with *L. major* and iron status in different organs were monitored systematically with the development of cutaneous lesion. At week 10 post infection when there was maximum lesion development in the parasite infected left hind footpad, the iron content increased significantly in this tissue with the concomitant increase in parasite burden. *L. major* infection mediated iron accumulation in infected mouse footpad was found to be due to transferrin receptor upregulation and natural resistance-associated macrophage protein 1 (Nramp1) downregulation. Surge in iron level was found to be associated with the reduced hepatic iron storage that resulted increased serum iron. Limited iron storage in liver and bone-marrow of infected mice caused reduced hemoglobin level and production of deformed erythrocytes. Interestingly, *L. major* infected mice developed splenomegaly with significant upregulation of erythroid related genes. Importantly, oral iron supplementation post infection rescued the development of cutaneous lesion in infected mice. Together, our study unravelled a comprehensive mechanism behind developing iron deficiency anemia during cutaneous leishmaniasis and a novel therapeutic route of treating this infection by delivering iron.

## Introduction

Leishmaniasis is a spectrum of human diseases caused by the protozoan parasites of the genus *Leishmania* that affects about 1 million people worldwide across 100 endemic countries (Burza et al., 2018). Depending on the species clinical manifestation of this disease varies from cutaneous lesion to visceral infections (Abadías-Granado et al., 2021; Kaye & Scott, 2011; Maltezou, 2010; Steverding, 2017). *Leishmania* parasites show a digenetic life cycle alternating between insect and mammalian hosts (Kaye & Scott, 2011). Infection starts when a female sandfly takes blood meal from a healthy individual and injects metacyclic promastigotes into the skin. Once inside the mammalian hosts, the parasites are rapidly engulfed by macrophage cells and upon internalization the pathogens are subsequently delivered to the phagolysosomes. Within phagolysosomal compartment *Leishmania* transforms into amastigotes and replicates till the cell bursts (Dostálová & Volf, 2012). Majority of the intracellular pathogens fail to survive within this nutrient limiting acidic compartment of macrophage cells which is packed of hydrolytic enzymes (Haas, 2007). However, genome wide transcriptome and proteomic analysis revealed *Leishmania* amastigotes are well adapted to function within phagolysosomes by maintaining their internal pH homeostasis and acquiring nutrients efficiently. Iron is a key micronutrient for all the organisms including *Leishmania* that acts as a cofactor in crucial enzymatic pathways (Cairo et al., 2006; Taylor & Kelly, 2010). Since iron level within macrophage phagolysosome is tightly regulated, *Leishmania* has been found to alter several host proteins involved in this process to maintain a continuous supply of this metal ion within their intracellular niche. For example, *Leishmania amazonensis (L. amazonensis)* has been found to promote fusion of transferrin bound endosome to parasite containing phagosomes to keep a continuous supply of iron to the parasites within infected macrophages (Luck & Mason, 2012). Additionally, in order to increase macrophage intracellular labile iron pool for their own benefit it causes downregulation of plasma membrane iron exporter, ferroportin (Fpn) (Ben-Othman et al., 2014). Recently, we have identified that infection of macrophages with *L. major* parasites inhibits phagolysosomal iron exporter, natural resistance associated macrophage protein 1 (Nramp1) expression which in turn helps in parasite growth by increasing phagolysosomal iron availability (Banerjee & Datta, 2020). Together, all these findings show how *Leishmania* infection severely impairs macrophage iron homeostasis. Macrophage cells are not only important in our defence system against invading pathogenic infection but also in regulating physiological iron homeostasis (Soares & Hamza, 2016). This is evident from the fact that impairment in its function leads to various clinical disorders including hypoferremia, anemia (Ganz, 2012). Although *Leishmania* infection is known to alter macrophage iron metabolism, very little is known on the effect of this alteration over systemic iron homeostasis. Also, how different species of *Leishmania* effects this process with the progression of disease is yet to be fully understood.

Among all the species, *Leishmania donovani (L. donovani)* is known to visceralize in an infected individual where the parasites infect and grows in phagocytes of the bone marrow, spleen and liver resulting into massive hepatosplenomegaly (Kaye & Scott, 2011; Maltezou, 2010). Detailed investigation revealed that *L. donovani* infection alters the erythropoiesis in the spleen and bone marrow by affecting erythroid progenitors by inducing a stress response on the expression of erythroid specific genes (Lafuse et al., 2013). The progressive replication of the parasites within these organs is perhaps thought to be associated with the ability of causing this hematopoietic alteration. However, manipulation of all these tightly regulated process in the reticuloendothelial organs leads to the severe form of anemia during visceral leishmaniasis which pose a serious health problem (Lafuse et al., 2013). Contrary to this, cutaneous leishmaniasis (CL) caused by *L. major* and its related species is the most common form of leishmaniasis that results into disfiguring skin lesions (Abadías-Granado et al., 2021; Steverding, 2017). Although lymphatic spread and lymph-gland involvement may precede lesion development the spread of infective parasites within internal organs is very uncommon. Also, inflammation and ulceration has only been reported at the site of infection (Palumbo, 2010; Zulfiqar et al., 2017). Although iron deficiency during CL has not gained a similar attention like visceral form of this disease previous report shows that children infected with cutaneous leishmaniasis suffers with reduced hemoglobin level with concomitant development of iron deficiency anemia compared to non-infected child groups (Weigel et al., 1995). In fact, children with past CL infection are prone to develop clinical symptoms associated with malnutrition. Furthermore, exogenously delivered iron to *L. major* infected mice at early stages of infection has been found to be effective in restricting the progression of this disease (Bisti et al., 2000). Together, all these indicate that CL infection leads to the development of iron deficiency in an unknown mechanism. In fact, early development of iron deficiency whether linked to host susceptibility towards CL progression due to reduced cell-mediated immunity involving defective neutrophil and macrophage killing function needs to be investigated. In this present study, we attempted to understand these unresolved issues on the development of iron deficiency anemia and its underlying mechanism during CL infection using the well-established *L. major*-Balb/c model system. We report here that *L. major* infection causes iron overload at the site of lesion development by altering the iron importer transferrin receptor 1 (TfR1) and iron recycling protein Nramp1 expression which is associated with increased parasite replication. Interestingly, our work shows that increased supply of this iron to the site of infection is coming through blood which is possibly due to reduced hepatic iron storage during infection. In fact, we found that *L. major* infection promotes reduced hemoglobin and bone marrow iron content that results into drastic alteration in normal erythrocyte population by inducing stress erythropoiesis response in spleen. Moreover, we report that cutaneous lesion development and its associated iron deficiency can be restored to normalcy by inducing an oxidative stress to the parasites via oral iron supplementation. Therefore, besides identifying the hitherto unknown mechanism of iron deficiency anemia during cutaneous leishmaniasis this study also establishes that despite colonizing to the site of infection *L. major* parasites are also able to alter systemic iron homeostasis involving reticuloendothelial organs.

## Results

### Iron is trafficked to the site of *L. major* infection

*L. major* infection is known to cause chronic cutaneous lesions in human as well as in experimental animals which is restricted to the site of infection. Iron being the essential micronutrient for the parasitic growth, we were curious to know the status of iron at the infection site. For this, we infected Balb/c mice with *L. major* promastigotes by sub-cutaneous injection to the left hind footpad and monitored the development of cutaneous lesion up to week 10 (Gurung et al., 2015). It is evident from Fig.1A and S1A that starting from week 4 p.i. the lesion started to develop and at week 10 p.i. we could detect a severe lesion in the infected footpad. To check if lesion development is correlated with parasite burden and how it affects the footpad tissue morphology, we next sought to quantitate the number of *Leishmania* parasites in the lesion site and examine the footpad tissue architecture during the course of infection. As documented in Fig. S1B, our histological investigation revealed that *L. major* infected mouse developed severe inflammation marked by increased accumulation of immune cells in the infected footpad (Gurung et al., 2015). Throughout these time points we could also identify increasing number of *Leishmania* amastigotes at the lesion site (Fig. S1C). We next checked iron concentration of in the mouse footpad during infection. For this, we harvested the left hind-footpad from both uninfected and *L. major* infected mouse and measured the iron content in the whole tissue lysates by ferrozine-based colorimetric assay (Fig. 1B). At week 4 p.i. the iron content within infected mice footpad was almost similar to uninfected mice footpad. However, at week 7 p.i. the iron level in infected mice footpad increased significantly (100μg/ mg tissue) compared to its uninfected counterpart (75μg/ mg tissue). Interestingly, at week 10 p.i., when there was a maximum swelling as well as parasite load, we observed a more than two folds increase in iron content of *L. major* infected mouse footpad (~ 150μg/ mg tissue) compared to its uninfected control (~70μg/ mg tissue) (Fig. 1B). These results unambiguously established that proliferation of *L. major* amastigotes is accompanied by flow of iron at the site of cutaneous lesion.

**Figure 1.**
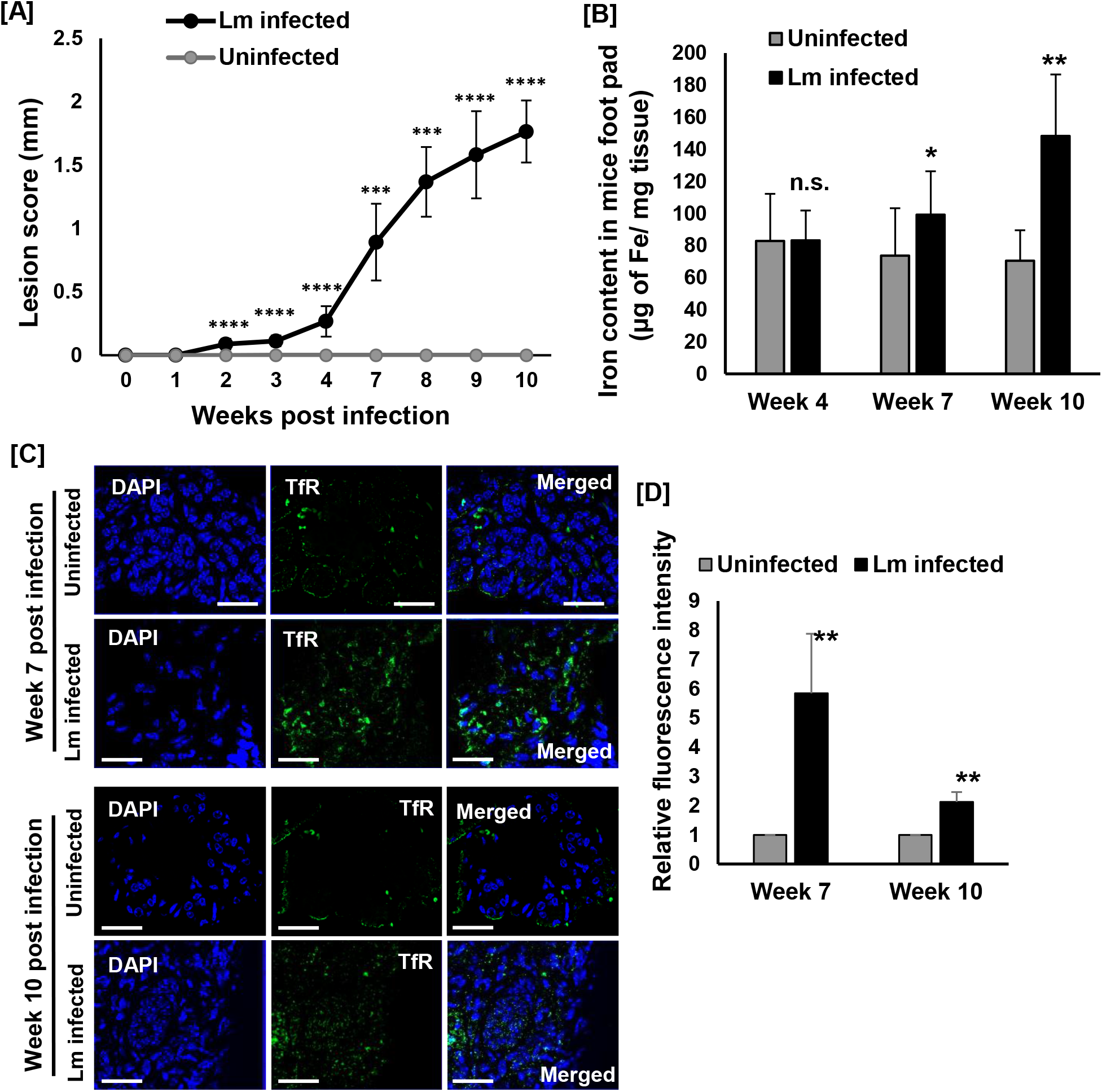
*L. major* infection causes iron overload in the mouse footpad by upregulating TfR protein expression. Balb/c mice were either uninfected or infected with 1×10^6^ stationary phase promastigotes of *L*. *major* parasite subcutaneously into the hind footpad. (A) Footpad swelling after *L*. *major* infection in each mouse was monitored by weekly measurement of width and thickness of the infected foot. The lesion score was measured upto week 10 p.i. as described earlier against the measurement obtained from uninfected footpad. Each group of this experiment contained at least 5 mice and repeated thrice. (B) At week 4, 7 and 10 post infection mice were euthanized and iron content was measured in both uninfected and Lm infected mice left footpad using ferrozine assay and represented in the bar diagram. Error bars represent SEM values calculated from three independent experiments with each experimental group having 5 animals. *p≤ 0.05; **p ≤ .01; ***p≤ 0.001; ****p ≤ .0001. (C) TfR protein level was visualized by immunostaining with anti-TfR (green) in uninfected or *L. major* (Lm) infected mouse footpad sections at the indicated time points (week 7 and 10 p.i.). Nuclei were stained with DAPI (blue). Tissue sections were visualized under 63X oil immersion objective of Zeiss Apotome microscope. Scale bar: 20μm. (D) Bar diagram representing the quantitative estimation of TfR expression as indicated by the relative fluorescence intensity of TfR at indicated time points in uninfected (grey bar) and Lm infected (black bar) mouse footpad sections. Several fields were imaged using the above mentioned microscope and images were analysed for the fluorescence intensity measurement using ZEN software of the microscope. Uninfected footpad sample were used as reference sample during quantification. Error bars represent SEM calculated from three independent experiments. *p≤ 0.05; **p ≤ .01; ***p≤ 0.001; ****p ≤ .0001; n.s., non-significant.

### Altered expression of TfR1 and Nramp1 in *L. major* infected mouse footpad

Transferrin receptor 1 (TfR1), the carrier protein of transferrin is known to import iron within the cells (Mackenzie et al., 2008; S. Wang et al., 2020). Importantly, *Leishmania* is known to upregulate the expression of TfR1 in macrophages in order to make iron available for their growth (Das et al., 2009). Hence, we wanted to check if increased accumulation of iron within mouse footpad is due to altered expression of TfR1 protein. Our immunofluorescence staining of uninfected and *L. major* infected mouse footpad tissue sections with anti-TfR1 revealed that compared to the uninfected mouse footpad the TfR1 protein expression was significantly upregulated (by about 5 folds and 2 folds at week 7 and 10 p.i., respectively) in *L. major* infected mouse footpad (Fig. 1C-D). In iron loaded cells, Nramp1 is known to play a crucial role in recycling of iron within tissues. Recently, we reported that *L. major* infection severely inhibits Nramp1 expression in macrophage cells which in turn results in increased iron accumulation within the phagolysosome that supported parasite growth (Banerjee & Datta, 2020). Hence, we next attempted to check the status of Nramp1 in mouse footpad at week 7-10 p.i. Our immunofluorescence staining of footpad sections using anti-Nramp1 shows distinct intracellular punctate appearance of this protein in uninfected tissue. In the infected mouse footpad, there was a drastic reduction (of more than 2 folds) in Nramp1 protein level at week 7 and 10 p.i. as compared to its uninfected counterpart. Thus, our in vivo results are consistent with our previously reported in vitro cell culture-based data showing that *L. major* infection suppresses Nramp1 protein expression (Fig. 2A-B). Collectively, our findings suggests that *L. major* infection is inducing TfR1 mediated iron uptake and simultaneously blocking its recycling via inhibiting Nramp1 expression in order to acquire the micronutrient at the site of infection. This further intrigued us to undertake an in-depth physiological investigation to understand the source of this elevated iron.

**Figure 2.**
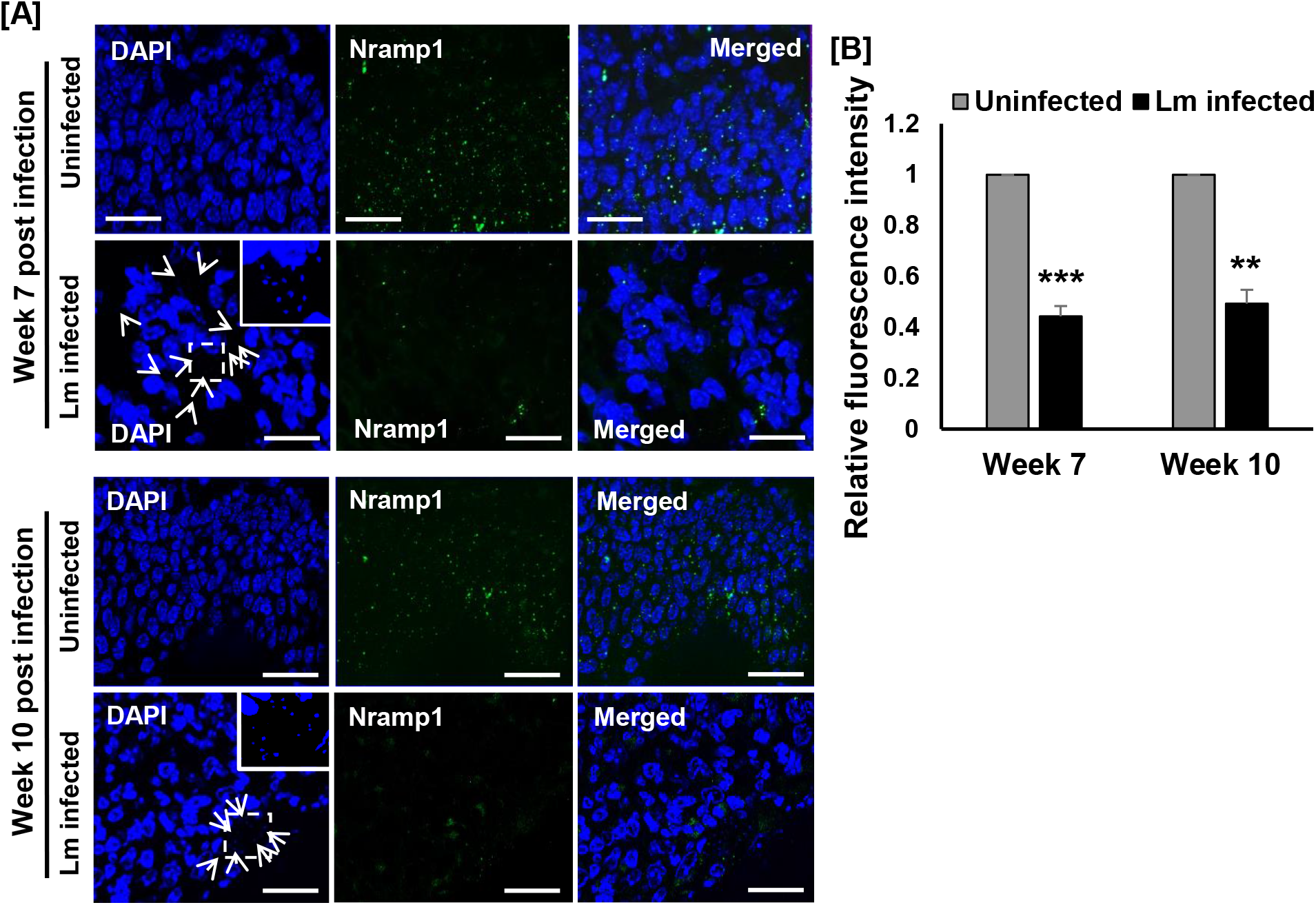
*L*. *major* infection promotes downregulation of Nramp1 at the site of infection in mice. At week 7 and 10 post infection, *L. major* infected mice were euthanized, (A) footpad tissue sections were fixed and further immunostained with anti-Nramp1 (Green) to check the protein level in both uninfected and infected mice footpad. DAPI (blue) was used to stain the nucleus. Zeiss Apotome microscope was used to visualize the tissue staining pattern with ×63 oil immersion objective. Scale bar: 20μm. Bar diagrams in (B) show the comparative analysis of relative fluorescence intensity of Nramp1 protein between uninfected and Lm-infected macrophages at both the time points quantified using ZEN software where uninfected cells has been the reference sample. Error bars represent SEM calculated from three independent experiments. n.s., non-significant; *p≤ 0.05; **p ≤ .01; ***p≤ 0.001; ****p ≤ .0001.

### *L. major* infection in the footpad resulted in increased serum iron, reduced hemoglobin and damageds RBCs

We next investigated whether rush of iron at the site of *L. major* infection in the footpad is associated with an increased level of serum Iron. The transferrin (Tf) bound iron circulates in the bloodstream, which is taken up by the cells of various tissues via transferrin receptor 1 mediated endocytosis (Luck & Mason, 2012). Therefore, we evaluated the serum iron level and the transferrin saturation in both uninfected as well as in *L. major* infected mice at 4-10 weeks post infection. Although at week 4 p.i. there was no significant difference between uninfected and infected mice, we observed a substantial increase in the serum iron level at week 7-10 p.i. in infected mice (Fig 3A). The transferrin saturation also showed a significant increase in the infected mice at week 7-10 p.i., confirming the presence of excess serum iron upon *L. major* infection (Fig. 3B). This intriguing results prompted us to examine the haemoglobin (Hb) level in both uninfected and infected mice blood which is as a potential indicator of physiological iron overload or deficient condition (Camaschella, 2017). Interestingly, while there was a significant increase in serum iron as well as transferrin saturation level at 7 and 10 weeks p.i., we observed a markedly reduced Hb in *L. major* infected mice blood as compared to uninfected mice (Fig. 3C). We also found that in infected mice at week 7 p.i. there was two folds reduction in bone marrow iron content and which further decreased by about 4.5 folds at week 10 p.i. as compared to the uninfected mice (Fig. S2A). Since Hb is primarily responsible for maintaining the structural as well as functional integrity of matured red blood cells (RBCs) we next sought to examine if there is any morphological alteration of RBCs of *L. major* infected mice groups. For this, we prepared blood films from both the uninfected and infected mice groups at week 7-10 p.i. and stained it with the Giemsa stain. Interestingly, our microscopic observation revealed that there was a major discrepancy in the overall shape of the RBCs in *L. major* infected mice (Fig. 3D). This led us to critically analyse and compare the morphological appearances of those erythrocytes with the previous reports. Although at week 10 p.i. we did not find the presence of cigar cells in infected mice blood smear, at both the time points (7and 10 p.i) we identified the presence of hypochromic, polychromatic RBCs, spherocytes, stomatocytes and dacrocytes (Fig. S2B) (Ford & Ford, 2013; Zabolotzky & Walker, 1990). Further, our quantitative analysis of these microscopic observations showed ~ 50% drop in the number of normocytic erythrocytes in *L. major* infected blood as compared to uninfected mice. (Fig. 3E). Collectively, the presence of these crippled erythrocytes in *L. major* infected mice indicates development of poikilocytosis at week 7-10 p.i. which is demonstrates a systemic iron deficient condition in these mice (Zabolotzky & Walker, 1990).

**Figure 3.**
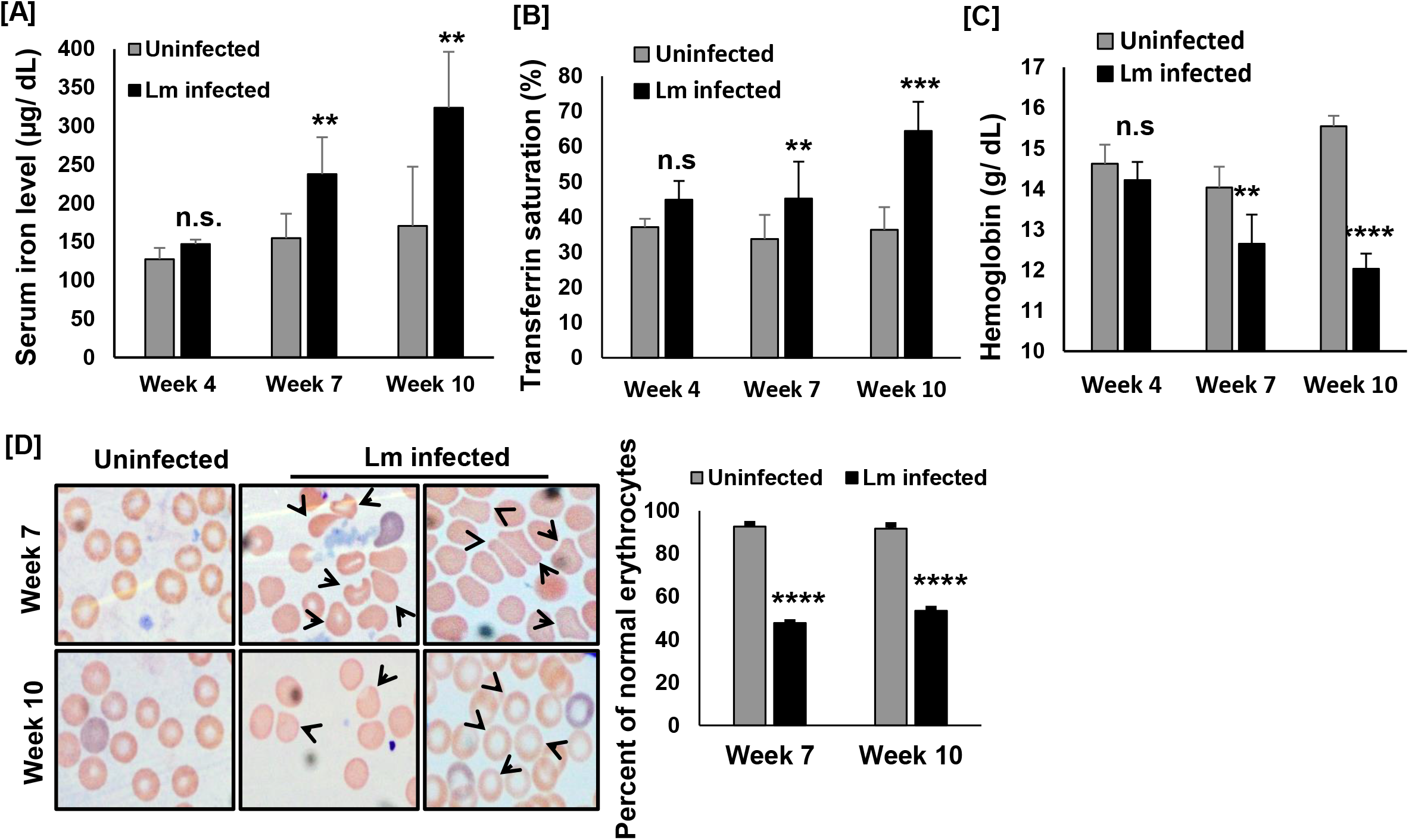
*L. major* infection in mice footpad promotes increased serum iron, reduced hemoglobin level and damaged RBCs. 6-8 weeks old female Balb/c mice were infected with *L. major* parasites were used to collect blood at week 4, 7 and 10 post infection along with from their uninfected control mice. (A) and (B) Further, serum these blood samples were used to prepare the serum and measured the iron content and transferrin saturation in it by using the tibc kit as mentioned in the materials and methods section. (C) The blood collected from those experimental mice were directly used to measure the hemoglobin level in it by using the Hemocor D solution. Results are represented with a bar diagram in a comparative manner between uninfected and Lm infected mice. (D) Blood smear was prepared and stained with Giemsa and further visualized under the microscope. Arrows (black) in the Lm infected panels showing the presence of altered RBCs by shape. Scale bar 10μm. Further, quantitative analysis of these microscopic images were performed to dissect the percent of normal erythrocytes present in both uninfected (grey bar) and Lm infected mice (black bar) blood at week 7 and 10 post infection and represented in the bar diagram. For this quantification, at least 2000 cells from each mice were counted. From both the experimental groups at least 5 mice were used to collect the blood from three independent experiments. Error bars represent SEM calculated from three independent experiments. n.s., non-significant; *p≤ 0.05; **p ≤ .01; ***p≤ 0.001; ****p ≤ .0001.

### *L. major* infection in the footpad results into hepatic iron deficiency

Since liver, the reservoir of iron, tightly maintains the availability of this micronutrient at the systemic level our results prompted us to check the status of liver iron storage in *L. major* infected mice. For this, we harvested livers from uninfected mice or mice infected with *L. major* in the footpad. Despite the localized infection in the footpad, strikingly, we noticed a discoloration of infected mice liver at week 10 p.i. as compared to the uninfected mice liver (Fig. 4A). We speculated that this discoloration of liver might be due to reduced hepatic iron content. However, we did not observe any difference in liver weight in between uninfected and infected mice suggesting *L. major* infection does not cause hepatomegaly (Fig. S3A) (Burza et al., 2018). Our quantitative ferrozine based assay of the liver lysates at week 7-10 p.i. revealed that at both the time points there was a drastic reduction (more than 2 folds) in the liver iron content of infected mice as compared to the uninfected mice. However, at week 10 p.i. the reduction in hepatic iron content of infected mice was more prominent (about 4 folds) (Fig. 4B). In this context, it is worth mentioning that our H&E staining of liver sections at week 7 and 10 p.i. showed that there was a severe loss of tissue integrity in *L. major* infected mice liver with granuloma formation and lymphocytic infiltration. Interestingly, we also noticed sinusoidal dilation in the infected mice liver at both these time-points (Fig. S3B). Ferroportin is known to be the sole iron exporter present in the cell membranes of hepatocytes (Donovan et al., 2005). Hence, to understand the mechanism behind reduced hepatic iron content we examined Fpn expression by qRT-PCR. Although at week 7 p.i. we observed a significant 3-fold upregulation of Fpn mRNA in *L. major* infected mice compared to its uninfected control mice, at week 10 p.i. there was a several fold reduction in Fpn expression (Fig. 4C). We did not observe any difference in iron importing TfR1 transcription level at week 7 p.i. in infected mice. But there was more than 2-fold reduction in TfR1 mRNA level at week 10 p.i. in infected mice compared to uninfected mice (Fig. 4D). We further checked the expression of ferritin in the liver (Anderson & Shah, 2013). Importantly, at both the time points post infection there was a drastic inhibition in Ferritin transcript level which further indicates reduced hepatic iron level during *L. major* infection (Fig. 4E). Together, these results suggest that increased iron export as well as its reduced import contributes to hepatic iron deficiency.

**Figure 4.**
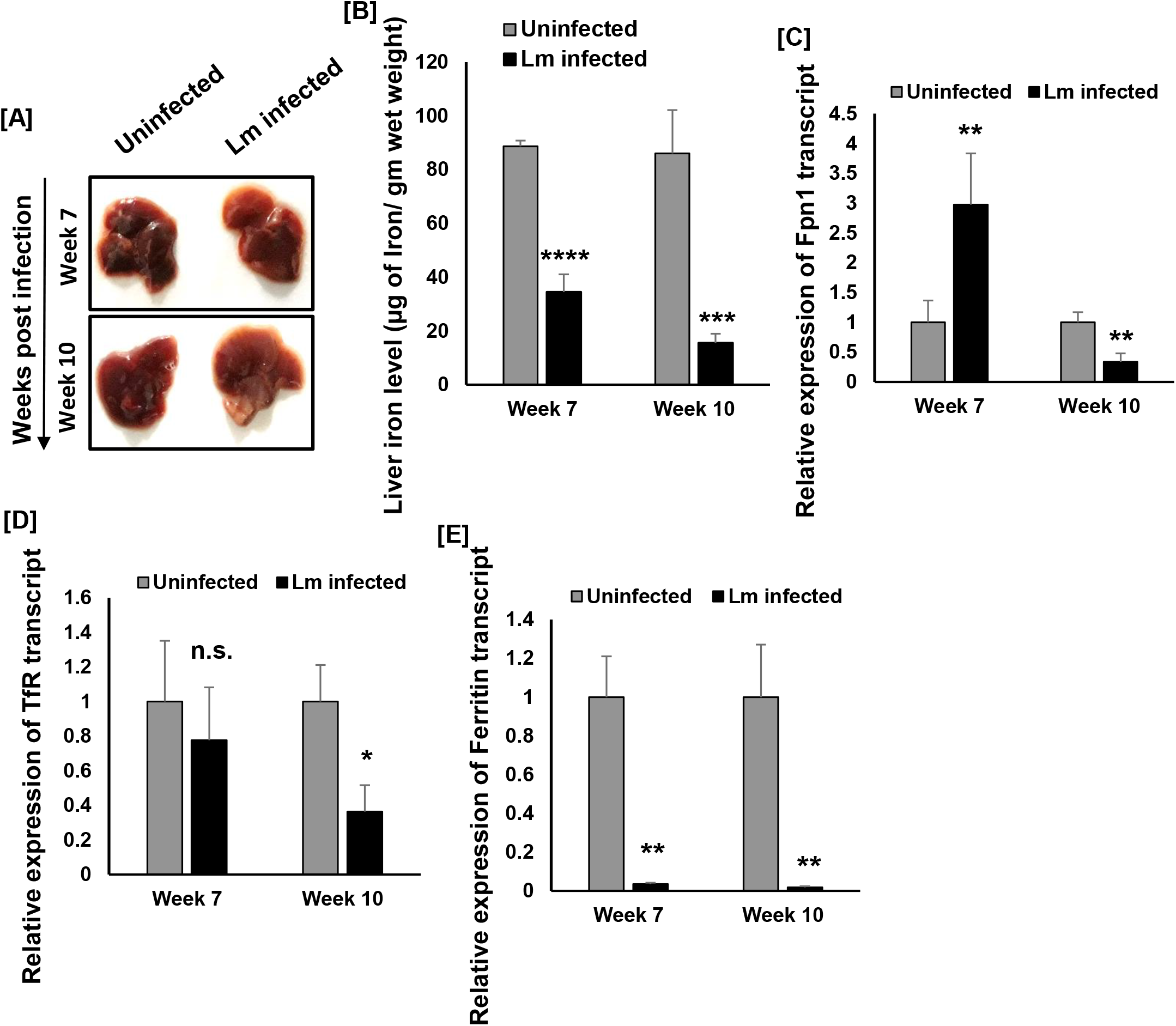
Hepatic iron deficiency is associated with the progression of cutaneous lesion. At week 7 and 10 post infection mice were euthanized and liver was observed for any alteration. (A) Representative image showing livers of both uninfected and Lm infected mouse where discolouration of infected mouse liver is visible. (B) Post harvesting the liver lysates were prepared and iron content in it was measured by ferrozine assay in both uninfected (grey bar) and Lm infected (black bar) mice groups which is represented in the bar diagram. Total RNA was prepared from both the uninfected and Lm infected mouse liver and further the cDNA was used to check the level of either Fpn1 (C), TfR (D) and ferritin (E) transcript level by qRT-PCR. The bar diagrams are representing the quantitative analysis of expression level of those genes between uninfected and Lm infected mice groups. Error bars represent SEM values calculated from three independent experiments with each experimental group having 5 animals. n.s., non-significant; *p≤ 0.05; **p ≤ .01; ***p≤ 0.001; ****p ≤ .0001.

### *L. major* infection causes splenomegaly and triggers extramedullary erythropoiesis

It has been reported that spleen, in partnership with liver, regulates the physiological iron homeostasis by mediating degradation of senescent RBCs and further recycling iron into circulation (Nemeth & Ganz, 2006; Theurl et al., 2016). This led us to measure the iron content in spleens in uninfected and *L. major* infected mice at week 7 and 10 p.i. Our qualitative representative images as well as the quantitative analysis did not show any difference in splenic iron content in between uninfected and *L. major* infected mice at week 7 or 10 p.i. (Fig. 5A and S4A). However, it is worth noting that there was an enlargement in the spleen of infected mice at week 10 p.i. Further analysis of splenic length and weight demonstrates that at this time point both the parameters have significantly increased in infected mice spleen as compared to uninfected mice leading to splenomegaly (Fig. S4B-C) (Melo et al., 2020). Our microscopic observation of hematoxylin eosin-stained spleen sections showed disorganized follicular regions in infected mice as compared to uninfected mice at both the time points. In fact, the red pulp and T-cell areas are also distorted upon infection as represented in Fig. 5B. While studying the splenic architecture we found the presence of a substantial number of megakaryocytes in infected mice (Fig. 5B). This observation was extremely important since accumulation of megakaryocytes in infected spleen clearly indicates the ongoing hematopoietic activity within infected mice spleen (Huang & Cantor, 2009). Although replication of megakaryocytes in spleen during *L. donovani* infection has been reported earlier it is unusual and interesting in case of *L. major* infection since the parasite does not reside in this organ. In line with this finding, we observed a significant ~3 folds and ~7 folds increase in erythroid related genes, Epo-R and GATA1 expression respectively in infected mice at week 7 p.i. (Fig. 5C-D). Although at week 10 p.i. GATA1 expression remained almost same we observed ~6 folds upregulation in Epo-R transcript level in infected mice spleen (Fig. 5C-D). We also inspected the status of globin chain synthesis regulatory genes, Sox6 and Klf1 (Dumitriu et al., 2010; Siatecka & Bieker, 2011). As shown in Fig. 5E-F at 7 weeks p.i. both these genes shown similar trend, where we observed a significant 2 folds upregulation in infected mice as compared to uninfected mice. However, at week 10 p.i. there was further upregulation where Sox6 expression increased about 5 folds and in case of Klf1 we observed ~ 4 folds upregulation in infected mice (Fig. 5E-F). Together, our results for the first time show that during localized *L. major* infection, iron content within spleen remains unaltered, however, the expression of erythroid related genes increased abruptly which has already been recognized as the stress erythropoiesis response in mice (Liao et al., 2018; H. Wang et al., 2021).

**Figure 5.**
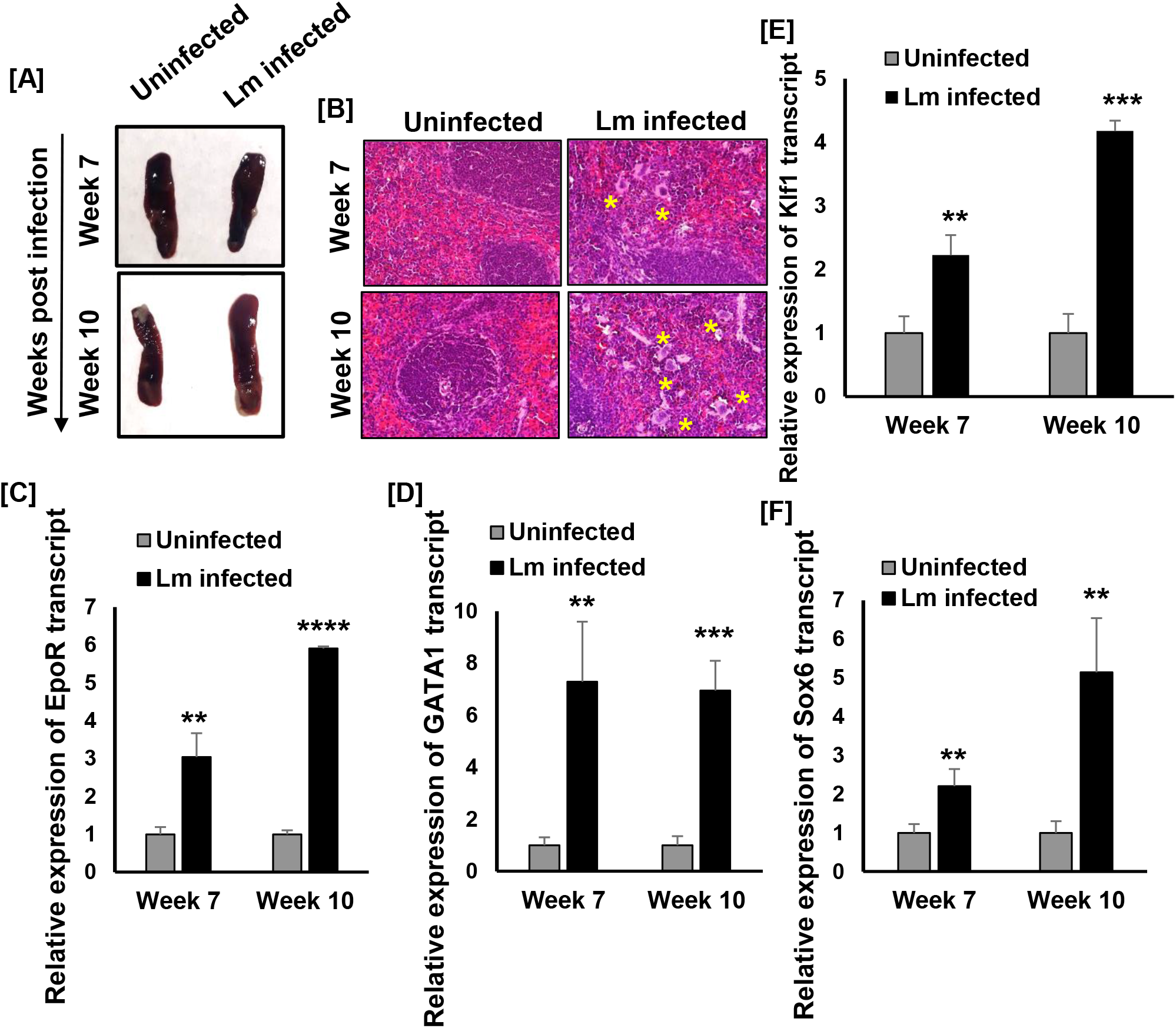
*L. major* infection causes splenomegaly and stress erythropoiesis. At week 7 and 10 post infection mice were euthanized and (A) both uninfected and infected mice spleen were harvested and the morphological alterations are represented in the figure. (B) spleen tissue sections were prepared and further stained with H&E to visualize the tissue architecture of both uninfected and *L. major* (Lm) infected spleen. In the Lm infected panel distortion of red pulp and white pulp regions are visible where stars are indicative of the presence of megakaryocytes. Scale bar 10μm. Total RNA was prepared from both the uninfected and Lm infected mouse spleen and further the cDNA was used to check the level of either EpoR (C), GATA1(D), Klf1 (E) and Sox6 (F) transcript by qRT-PCR. The bar diagrams are representing the quantitative analysis of expression level of those genes between uninfected and Lm infected mice groups. Error bars represent SEM values calculated from three independent experiments with each experimental group having at least 5 animals. n.s., non-significant; *p≤ 0.05; **p ≤ .01; ***p≤ 0.001; ****p ≤ .0001.

### Oral supplementation of iron restricts cutaneous lesion development in *L. major* infected mice

Since our experimental results clearly shows *L. major* infection promotes iron deficiency, we next sought to examine whether iron supplementation can restrict parasitic infection. For this, we first infected the J774A.1 macrophage cells as well as thioglycolate-elicited peritoneal macrophages from Balb/c mice with *L. major* promastigotes and observed the impact of iron supplementation on parasite growth. As shown in Fig. S5A-B, at 12hrs post infection, 100μM ferric ammonium citrate (FAC) treated macrophages indeed had significantly lower level (more than 2 folds) of intracellular parasite burden (amastigotes/ macrophage). Following this lead, we were interested to check the kinetics of disease progression in *L. major* infected mice upon FAC treatment. Previously, it has been shown that *L. major* infection related lesion development can be treated with iron supplementation through intra-peritoneal injection (Bisti et al., 2000). We performed infection studies in Balb/c mice with *L. major* promastigotes as described earlier in this manuscript. Apart from the uninfected and *L. major* infected mice groups, one group of mice were infected with *L. major* promastigotes and treated with 2mg/ kg of FAC via oral gavage 4 times in a week throughout the experiment and another group of control mice (uninfected) only received the iron supplementation (Kulda et al., 1999). All these mice were examined weekly for the disease development and the cutaneous lesion were measured as described earlier. Although *L. major* infected mice developed uncontrolled footpad lesion upto 10 weeks post infection as observed by us in our previous experiment there was almost a complete recovery of lesion development associated with a drastic reduction in parasite burden in mice which received oral iron supplement (Fig. 6A-B). Collectively, our data for the first time shown that oral iron supplementation is highly effective in treating cutaneous lesion development during *L. major* infection.

**Figure 6.**
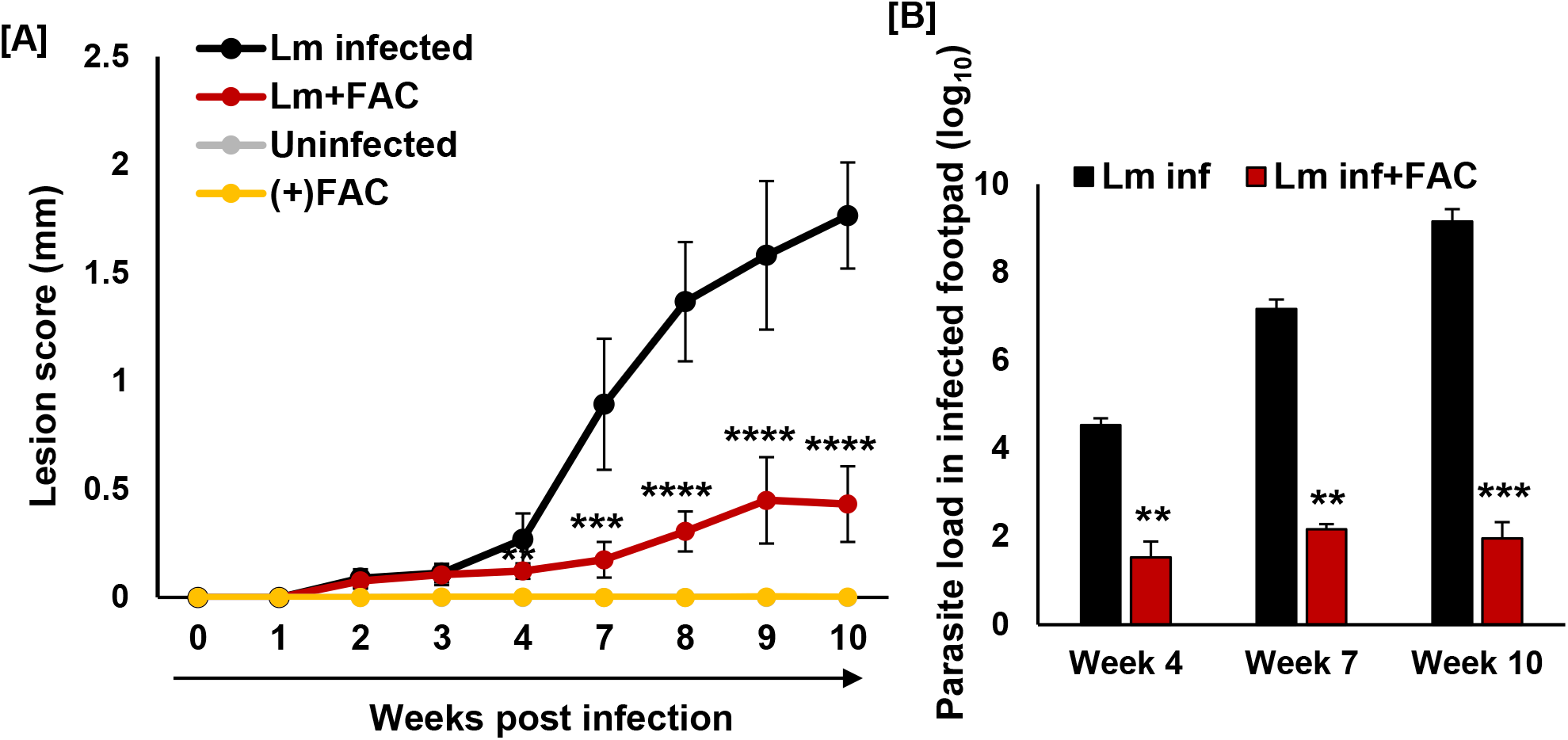
Iron supplementation inhibits *L. major* infection. 6-8 weeks old female Balb/c mice were anaesthetized and 1X10^6^ *L. major* promastigotes were injected into left hind footpad and lesion development was observed upto 10 weeks post infection (p.i.) in uninfected (grey), Lm infected (black), ferric ammonium citrate (FAC) (yellow) supplemented or Lm infected and FAC supplemented (red) mice. (A) Line diagram showing that lesion development is significantly stalled in FAC supplemented Lm infected mice. (B) Parasite load in the *L. major* infected mice footpad was determined at week 4, 7 and 10 post infection by limiting dilution assay which is represented in the bar diagram. Error bars represent SEM values calculated from three independent experiments with each experimental group having at least 5 animals. n.s., non-significant; *p≤ 0.05; **p ≤ .01; ***p≤ 0.001; ****p ≤ .0001.

### Oral iron supplementation promotes surge in ROS level at the site of *L. major* infection

Iron supplementation therapy has previously been found to be associated with increased susceptibility towards related pathogenic infections including *Salmonella* and *Mycobacteria* (Nairz et al., 2014). Since our results demonstrated a contrasting outcome, we were intrigued to explore the mechanism how iron supplementation could effectively restrict cutaneous lesion development in *L. major* infected mice. For this, we first harvested the footpad tissues from different groups of mice at 7-10 weeks post infection and measured the iron content in them by ferrozine-based assay. Interestingly, compared to *L. major* infected mice, the footpad iron content remained significantly high (~1.5 folds) in *L. major* infected that were fed with FAC (Fig. 7A). This observation indicates that oral iron supplementation promotes accumulation of excess iron to the *L. major* infected mice footpad tissue. Whether this excess iron causes restriction of parasite growth by increased burst of reactive oxygen species (ROS) was our next question (Zhou et al., 2018). Hence, we quantified ROS status using the fluorescence dye DCFDA based detection method by fluorescent activated cell sorting (FACS) technique (Eruslanov, E., 2010). As shown in Fig. S6A-B, as compared to uninfected peritoneal macrophages there was about 4 folds increase in cellular ROS level upon infection to *L. major* at 12hrs post infection. However, it is extremely important to note that the ROS production was further elevated by ~2 folds in *L. major* infected FAC treated macrophages compared to the infected control (Fig. S6B). Nevertheless, our results certainly confirms that inhibition of parasite growth during iron treatment is potentially occurring due to the toxic effect of excess oxidative burst in those peritoneal macrophages. Hence, our observation of excess iron accumulation in *L. major* infected mice footpad upon oral iron feeding at week 10 p.i. further prompted us to check the status of ROS in this tissue. For this, we harvested the footpad tissues from those mice groups and prepared the lysate using mild treatment of collagenase. Further, the debris were removed and the ROS level in those footpads cell suspension was measured by FACS using the fluorescent dye DCFDA. Interestingly, in *L. major* infected mice we have observed about ~2.5 folds increase in ROS level as compared to uninfected mice. However, it is worth mentioning that in *L. major* infected FAC fed mice the footpad ROS level is much higher, which was ~4 folds as compared to uninfected mice and also ~2 folds more compared to its infected control (Fig. 7B-C). Together, these results clearly shows that oral iron supplementation of FAC promotes excess iron accumulation at the site of *L. major* infection that in turn potentially suppresses the lesion development via generating excess oxidative burst. However, we wanted to check if it causes any iron overloaded condition in the major iron storage organ, liver of those animals. From Fig. S7A it is evident that at week 7-10 p.i. when the liver iron content has dropped several folds in control infected mice oral iron treatment helped to recover it in the infected mice group. Since, we have already observed that reduced liver iron in *L. major* infected mice is associated with excess serum iron this important outcome further intrigued us to monitor the serum iron level in *L. major* infected iron fed mice. Interestingly, as shown in Fig. S7 B iron supplementation also brought back the serum iron level to normalcy. Together, these observations strongly suggest that oral doses of iron can rescue the cutaneous lesion as well as the systemic iron deficiency caused by *L. major* infection in experimental mice.

**Figure 7.**
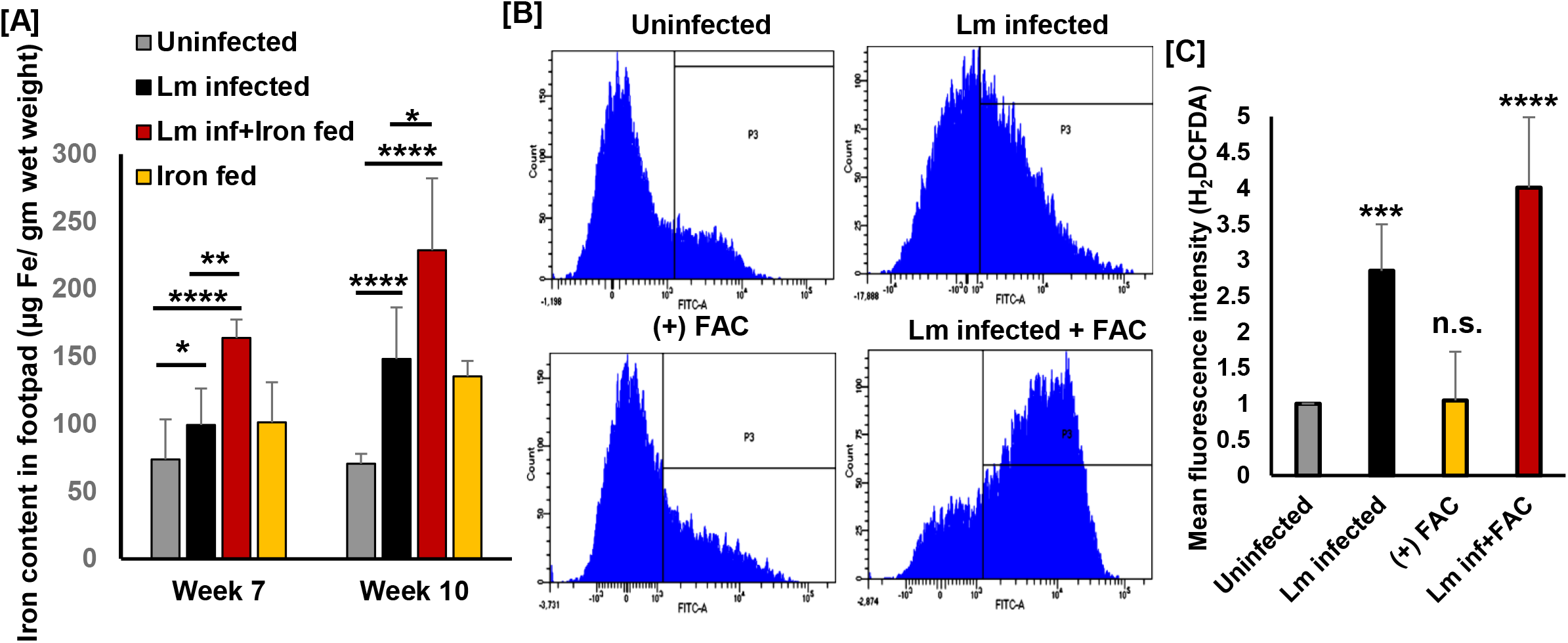
Iron supplementation causes surge in iron and ROS level in infected mouse footpad. At week 7 and 10 post infection mice were euthanized and footpad tissue were harvested from uninfected, Lm infected, FAC supplemented and Lm infected as well as FAC supplemented mice groups. (A) Further the lysates were prepared and iron content in this tissue was measured using ferrozine assay which is represented in the bar diagram. (B) Footpad tissue harvested similarly from those mice groups were treated with collagenase to release homogenous suspension of cells and further ROS level was measured using DCFDA based FACS assay. (C) Bar diagram is sowing the quantitative analysis of ROS level in those mice groups. Error bars represent SEM values calculated from three independent experiments with each experimental group having 5 animals. n.s., non-significant; *p≤ 0.05; **p ≤ .01; ***p≤ 0.001; ****p ≤ .0001.

## Discussion

In this study, we have shown that *L. major* infection severely alters the iron homeostasis at the site of infection as well as at the systemic level. Iron is a key micronutrient for all the organisms including *Leishmania* that acts as a cofactor in crucial enzymatic pathways. Although *L. major* infection is known to alter macrophage iron metabolism, very little is known on the effect of this alteration on systemic iron homeostasis. Since we have recently found that *L. major* infection promotes downregulation of macrophage Nramp1 it becomes crucial to identify what happens to this protein at the systemic level (Banerjee & Datta, 2020). Hence, we used the well-established Balb/c *L. major* infection model system to address this question. As a first step towards this, we measured the iron content in infected mouse footpad. Interestingly, at week 7 post infection when there was a significant lesion development in the footpad of infected mice, we found a heightened iron level in this tissue. It is also worth mentioning that at week 10 p.i. when there was maximum swelling of cutaneous lesion, the iron content was almost doubled in infected mice footpad as compared to uninfected mouse footpad. This observation was highly intriguing for us. Although *Leishmania amazonensis (L. amazonensis)* infection known to promote fusion of transferrin bound endosome to parasite containing phagosomes to keep a continuous supply of iron to the parasites within infected macrophages there is no such observations available at the tissue level within infected experimental animals (Luck & Mason, 2012). Since, transferrin receptors are the sole iron importer within cells or tissues, based on our observation we targeted to check if the increased iron accumulation is due to heightened expression of TfR. In line to our hypothesis, we found that while there was an increased iron level at week 7 and 10 p.i., TfR level was significantly upregulated. However, this might not be the only reason of this metal ion accumulation. Because, at the systemic level, alteration in iron accumulation may also be a result of disrupted recycling process (Beutler, 2007). Nramp1 is known to be the major iron recycling protein which mediates the release of processed iron from different complexes including hemoglobin. In this context, it is important to note that Nramp1 knockout mice develops severe iron accumulation within spleen indicating the fact that absence of this protein activity may lead to tissue iron deposition (Soe-Lin et al., 2009). Hence, taking the clue from our previous study, we also investigated if the heightened accumulation of iron is due to downregulation of Nramp1 at the infection site or not. For this, we attempted to check the status of Nramp1 within infected mice footpad for the first time using immunostaining. As revealed by our studies at both the time points including week 7 and 10 p.i. when there was significant iron accumulation, we failed to detect presence of Nramp1 in infected mice footpad tissue. We have already established that Nramp1 is the determining factor of *L. major* infection within macrophage cells. Interestingly, *in vivo* we also observed that while there was loss of Nramp1 protein at the infection site, the parasite burden was significantly high which is certainly supported by the increased availability of iron. However, it would be extremely rewarding to identify the source of iron flux to the site of infection and the status of systemic iron homeostasis under this situation. Although children infected with cutaneous leishmaniasis has shown reduced hemoglobin level with concomitant development of iron deficiency anemia the underlying mechanism has not been studied yet (Weigel et al., 1995). In fact, early development of iron deficiency whether linked to host susceptibility towards CL progression due to reduced cell-mediated immunity involving defective neutrophil and macrophage killing function needs to be investigated. In context to this, it is worth mentioning that previous report suggest that Nramp1 knockout mice develops severe iron accumulation within spleen and liver during erythrophagocytosis and haemolytic anemia which supported the notion that Nramp1 plays a critical role in efficient hemoglobin iron recycling within macrophage cells. Importantly, we have also observed that during *L. major* infection there is severe loss of Nramp1 within cutaneous lesion of infected mice that results into heightened iron accumulation. However, whether loss of Nramp1 during localized infection has any impact over systemic iron homeostasis or not needs to be checked. In fact, how the altered iron recycling within footpad tissue is linked to the surge in iron supply at that site is somewhat puzzling. Hence, we attempted to understand these unresolved issues on the development of iron deficiency anemia and its underlying mechanism during CL infection using the well-established *L. major*-Balb/c model system. Interestingly, our work shown that increased supply of this iron to the site of infection is mostly supplied through blood which is possibly due to reduced hepatic iron storage during infection. Initially, our blood related experiments shown that with the progression of lesion development and parasite burden with infected mouse footpad, the serum iron content and the transferrin saturation is increased. This observation was very interesting which definitely indicates that increased iron in infected mice footpad is supplied through blood. In this context, it is worth mentioning that *L. donovani* infection is known to deplete serum iron level while residing within reticuloendothelial organs (Lafuse et al., 2013). Hence, our observation certainly denotes a different mechanism in case of *L. major* infection. Importantly, we further found that at the late-stage post infection, the hemoglobin level is dropped significantly in infected mouse which was an outcome of reduced bone marrow iron content. Interestingly, it resulted into the alteration in erythrocyte morphology which gave us an indication that iron deficiency anemia might be prevailing in those infected animals. However, the most interesting observation was the iron deficiency in infected mouse liver. It is important to note that liver is the major iron storing organ of physiological system (Upton et al., 2003). Therefore, it is highly possible that in infected animals while the bone marrow iron level is reduced and anemic situation is prevailing, liver is responding to it by releasing its stored iron. This becomes more evident from our observation that expression of plasma membrane iron exporter Fpn is upregulated at week 7 p.i. in infected mice whereas the iron storing ferritin expression is almost negligible as compared to uninfected mice. Although we did not observe any difference in TfR transcript level at week 7 p.i. within infected mice liver, at week 10 p.i. there was a significant drop in the same. Since at week 10 p.i. the iron content in the infected mice liver is significantly low as compared to uninfected mice liver we speculate that it might be a host’s response to restrict further loss of iron by downregulating TfR expression. However, if the drop in iron level is linked to the altered hepatic tissue morphology remains an open question in case of cutaneous leishmaniasis. Since we wanted to identify the source of increased iron at the site of infection within infected mice, it was necessary to study the status of spleen. However, it was of a surprise to observe splenic enlargement in *L. major* infected mice as spleen enlargement is a well-known phenomenon that occurs during visceral leishmaniasis. This observation led us to systematically check the tissue morphology as well as the status of iron in it. Although we did not find any difference in iron content of spleen between uninfected and *L. major* infected mice our histopathological staining revealed the presence of megakaryocytes within infected mouse spleen. We speculated under the background of reduced hepatic iron storage and overall systemic anemic condition, accumulation of megakaryocytes would be a stress erythropoiesis response. It also becomes evident because splenic iron content remained unchanged throughout the time points post infection whereas the erythroid related gene expression including EpoR, GATA1 and Klf1 and Sox6 remained to be several folds high in infected mouse spleen as compared to its uninfected counterpart. Previously it has been found that stress erythropoiesis from spleen results into immature erythrocytes production (Liao et al., 2018; H. Wang et al., 2021). In line to these findings, our observations further confirms that the altered erythrocyte production during *L. major* infection is potentially linked to the stress erythropoiesis in spleen. Importantly, our data provides a new insight in cutaneous leishmaniasis where we are showing that restricted pathophysiology at the site of infection is causing altered systemic iron homeostasis. However, we cannot rule out the fact that loss of Nramp1 at the site of infection may induce different inflammatory signal in terms of producing cytokines which is involved in this overall malfunctioning. Therefore, it remained to be an interesting area of future research. But in this context the most promising outcome of our studies came from the oral iron supplementation experiment to the *L. major* infected mice. Although there is a report showing iron supplementation can restrict cutaneous leishmaniasis the approach was to deliver the excess iron though peritoneal injection (Bisti et al., 2000). Here, we have shown it more efficiently and clinically favoured route which is to provide orally. Our iron supplemented *L. major* infected mice had successfully inhibited the parasite growth by producing ROS at the site of infection. In fact, it also rescued the development of iron deficiency anemia in infected Balb/c mice. Collectively, all our findings elucidate a novel aspect of altered systemic iron homeostasis during cutaneous leishmaniasis where loss of Nramp1 at the site of infection is linked to the reduced hepatic iron storage and stress erythropoiesis by an yet to be identified mechanism which in turn mediates surge in iron supply to the lesion site. Moreover, we report that cutaneous lesion development and its associated iron deficiency can be restored to normalcy by inducing an oxidative stress to the parasites via oral iron supplementation. Therefore, besides identifying the hitherto unknown mechanism of iron deficiency anemia during cutaneous leishmaniasis this study also establishes that despite colonizing to the site of infection only *L. major* parasites are able alter systemic iron homeostasis involving reticuloendothelial organs. However, it would be extremely rewarding to check how the parasite is manipulating this process. Leishmania parasites are known to secrete a wide variety of biomolecules which alters host signaling pathways. Hence, it becomes essential to identify if the parasite is secreting any pathogenic factor while residing within the footpad in order to disrupt the balance between liver and spleen in regulating iron metabolism.

## Experimental Procedures

All reagents were purchased from Sigma-Aldrich unless mentioned specifically. Primers for PCR were obtained from Integrated DNA Technologies.

### *Leishmania* and mammalian Cell culture

The *L. major* strain 5ASKH was a kind gift of Dr. Subrata Adak (IICB, Kolkata). Briefly, *L. major* promastigotes were cultured in M199 medium (Gibco) pH 7.2, supplemented with 15% heat-inactivated fetal bovine serum (FBS, Gibco), 23.5 mM HEPES, 0.2mM adenine, 150 μg/ml folic acid, 10 μg/ml hemin, 120 U/ml penicillin, 120 μg/ml streptomycin, and 60 μg/ml gentamicin at 26°C. J774A.1 murine macrophage cells (obtained from National Center for Cell Sciences, Pune) were grown in Dulbecco’s modified Eagle’s medium (DMEM, Gibco) pH 7.4 supplemented with 2 mM L-glutamine, 100 U/ml penicillin, 100 μg/ml streptomycin, and 10% heat-inactivated FBS at 37°C in a humidified atmosphere containing 5% CO2. Cell number was quantified using a hemocytometer as reported by us previously (Banerjee & Datta, 2020).

### Peritoneal macrophage isolation from BALB/c mice

BALB/c mice obtained from the National Institute of Nutrition (NIN), Hyderabad, were housed in our institutional animal facility and used for experiments following the CPCSEA guidelines and Institutional Animal Ethics Committee approved protocol. Thioglycolate elicited peritoneal macrophages were isolated from 6-8 weeks old mice as described earlier (Banerjee & Datta, 2020). Briefly, 3% Brewer’s thioglycolate medium (Himedia) was injected into the mice peritoneum cavity and 4 days post injection, mice were euthanized and peritoneal macrophages were collected using 20 G needle. The isolated macrophages were then cultured in DMEM supplemented with 2 mM L-glutamine, 100 U/ml penicillin, 100 μg/ml streptomycin, and 10% heat-inactivated FBS, pH 7.4 at 37°C in a humidified atmosphere containing 5% CO2. After 18-24 h non-adherent cells were discarded and trypan blue dye exclusion test was performed to check cellular viability.

### Infection of macrophage cells with *L. major*

Macrophage cells were infected as described by us previously (Banerjee & Datta, 2020). Briefly, J774A.1 murine macrophages activated with LPS infected with stationary phase *L. major* promastigotes. Similarly, Balb/c isolated primary peritoneal macrophages were infected following identical procedure without activating them with LPS. The infection was allowed to continue for 12 hrs.

Following infection, cells were washed, fixed with acetone-methanol (1:1) and mounted with anti-fade mounting medium containing DAPI (VectaShield from Vector Laboratories) to visualize the macrophage and parasite nuclei. Quantification of intracellular parasite burden (number of amastigotes/100 macrophages) was performed by counting the total number of DAPI-stained nuclei of macrophages and *L. major* amastigotes in a field (at least 100 macrophages were counted from triplicate experiments).

### Infection of mouse with *Leishmania* parasites

All the animal experiments were performed on BALB/c mice following the ethical guidelines and protocol approved by the Institutional Animal Ethics Committee (IAEC) at IISER Kolkata. Infection of the experimental animals with *L. major* and subsequent lesion development in footpads was estimated as described previously with few modifications (Sacks & Melby, 2015). Briefly, 7-9 weeks old female BALB/c mice were anesthetized with an inhalation isoflurane, in a bell jar or gas anesthesia induction chamber and then infected subcutaneously with 1x 10^6^ late stationary phase wild type promastigotes in their left hind footpads (5mice/group in each of the experimental set a time). Lesion development in each mouse was monitored by weekly measurement of width and thickness of the infected foot. Similar measurements were taken from uninfected footpad of the same mouse as control using a digital caliper for a period of upto 10 weeks. Progression of disease is indicated by ‘lesion score’ which is estimated using the formula: [(width of infected footpad - width of non-infected footpad) x (thickness of infected footpad - thickness of non-infected footpad)]. During specific intervals of this animal infection studies parasite load in infected mice footpads were determined by limiting dilution assay (Bose et al., 2012; Sacks & Melby, 2015). Briefly, footpads were excised from the hind limbs and then sequentially washed in Wescodyne solution, 70% ethanol and finally in autoclaved water. Subsequently, excised tissue was weighed and then homogenized in complete M199 medium with the help of a micropestle. Next, each tissue homogenate was serially diluted in complete M199 medium and seeded in the same medium in a 24-well cell culture plate. The number of viable parasites per milligram of tissue was estimated from the highest dilution at which parasite could be grown after a period of 7-10 days incubation at 26°C.

### Antibodies and primers

Rabbit polyclonal anti-Nramp1 used in this study was developed by us as reported previously (ref). The anti-TfR antibody was kind gift of Dr. Arnab Gupta (IISER Kolkata). Total RNA was isolated from respective tissues of both uninfected and infected mouse using TRIzol reagent (Invitrogen). These were further treated with DNaseI (Invitrogen) to remove any DNA contaminations. The cDNA was synthesized using Verso cDNA synthesis kit (Thermo) from 1μg of total RNA. Transcript level of different genes was quantified using the following primers: GATA1-Forward 5’-CACTCCCCAGTCTTTCAGGTGTA-3’, GATA1 Reverse 5’-GGTGAGCCCCCAGGAATT-3’; Sox6 forward 5’-TTGGGGAGTACAAGCAACTGATGC-3’; Sox6 reverse 5’-ATCTGAGGTGATGGTGTGGTCGTT-3’; Klf1 Forward 5’-CCTCCATCAGTACACTCACC-3’; Klf1 reverse 5’-CCTCCGATTTCAGACTCACG-3’; TfR11 forward 5’-CAGAAAGTTCCTCAGCTCAACCA-3’; TfR1 reverse 5’-GTTCAATTCAACGTCATGGGTAAG-3’; Fpn1 forward 5’-CTACCATTAGAAGGATTGACCAGCT-3’; Fpn1 reverse 5’-ACTGGAGAACCAAATGTCATAATCTG-3’; ferritin heavy chain 1 forward GCGAGGTGGCCGAATCT; ferritin heavy chain 1 reverse 5’-CAGCCCGCTCTCCCAGT-3’; EpoR forward 5’-AAACTCAGGGTGCCCCTCTGGCCT; EpoR reverse 5’-GATGCGGTGATAGCGAGGAGAACC-3’; γ-Actin Forward 5’-GGCTGTATTCCCCTCCATCG-3’, beta-Actin Reverse 5’-CCAGTTGGTAACAATGCCATG T-3’. Real time PCR was performed using 7500 real time PCR system of applied Biosystems with SYBR green fluorophore (BioRad). Transcript levels of all the target genes were normalized with respect to γ-Actin expression in each of the sample.

### Estimation of tissue iron

To measure iron content within different tissue samples biochemical ferrozine assay was used as reported earlier (Riemer et al., 2004). Briefly, 20 μL of whole tissue lysates of either footpad or reticuloendothelial organs including liver, spleen and bone marrow was incubated with 100μl 10 mM HCl, 4.5% KMnO4 for 2 hrs at 60°C. The samples were then cooled down at room temperature. Further, the samples were incubated with freshly prepared iron detection reagent (6.5 mM of ferrozine, 6.5 mM of neocuproine, 2.5 M of ammonium acetate and 1 M of ascorbic acid) for 30mins. The absorbance of these solutions were measured at 550 nm using microplate reader. Iron content within these samples was then measured against a standard curve prepared using varying concentration of FeCl_3_ (0–300μM) as determined by us previously (Banerjee & Datta, 2020; Riemer et al., 2004).

### Haemoglobin measurement

To measure the blood hemoglobin (Hb) level reagents from the Coral Clinical systems was used which is supplied as HEMOCOR-D. In this assay Hb present in the blood is converted to methaemoglobin by reacting with potassium ferricyanide. Methaemoglobin further reacts with potassium cyanide to form cyanmethaemoglobin complex. Importantly, the measurement is based on a colorimetric technique where the absorbance of the product formed is measured at 540 nm. The intensity of the complex formed is directly proportional to the amount of Hb present in the sample. To check the amount of Hb present in both uninfected and *L. major* infected mouse at different time points post infection blood is first withdrawn from retro-orbital plexus using capillary. The blood was collected in EDTA solution to avoid the blood clotting. Following this 5 mL of Hemocor D solution was taken in test tube and 20μL of blood sample was mixed to it and further kept at room temperature for 5min. Next, the Hb content was measured by spectrophotometric measurement.

### Transferrin saturation and serum iron measurement

To measure the blood transferrin saturation level and serum iron content reagents from the Coral Clinical systems was used and the values were determined as per manufacturer’s protocol. Briefly, to measure the serum iron content, iron bound to transferrin is first released into the acidic medium and ferric iron is reduced to ferrous iron. The Fe^2+^ ion then reacts with ferrozine to form a violet coloured complex where the intensity of the colour is directly proportional to the amount of iron present in the sample quantified from the absorbance at 570nm. To measure the transferrin saturation following the manufacturer’s protocol, serum is first treated with excess amount of ferrous iron to saturate the iron binding sites of transferrin. Further, the excess iron is adsorbed and precipitated and the iron content in supernatant is quantified to detect total iron binding capacity (TIBC) using the specific solutions provided in the kit. Following this, transferrin saturation of the sample is calculated: Transferrin saturation= (serum iron/ TIBC) ×100.

### Giemsa staining of blood smear

A drop of blood collected from experimental mouse is taken at one end of a slide. Further the drop was spread using another slide holding at a 45° angle and backing into the drop of blood smoothly. Following this the slides were air dried and further fixed in methanol for 5-7 min. Slides were then air dried and stained with Giemsa solution (diluted in deionized water at a ratio of 1: 20). Following incubation for 60min in Giemsa stain solution the slides were taken out and rinsed in deionized water. Next, these were air dried and evaluated for the morphology of red blood cells and other corpuscles using inverted light microscopy at 40X magnification.

### Histopathology

First, anesthetized mouse is pin down with ventral surface facing up. Following wiping the surface area with 70% ethanol an incision is performed at the lower abdomen to open the viscera upto diaphragm so that heart and lungs will be exposed. Now the tip of the butterfly needle is inserted carefully in the left ventricle and perfusion with phosphate buffered saline is started. Immediately following this the superior and inferior vena cava is cut and perfusion is continued until liver turns pale yellow in colour. Now the needle is removed from PBS syringe and inserted into 4% paraformaldehyde containing syringe. Post perfusion with PFA the harvested organs were again stored into 4% PFA for 24hrs following which it was replaced with 70% ethanol. Harvested tissues were either stored long term or processed in the following way as per previously described methods (Das Sarma et al., 2000; Navas et al., 2001).

### Tissue processing

Fixed tissues were collected in tissue cassettes arranged with cassette holder and then placed into the automated tissue processor Citadel 2000. It processed the tissues kept in 70% ethanol with different treatments in the following order: first in 80% ethanol for 1hr, then in 90% and 95% ethanol for 1hr each. Further three consecutive treatments were performed in 100% ethanol for 1hr each. Next, the tissues were immerged in fresh xylene for 45 min three times. Finally, the tissues were treated with fresh paraffin two times for 1hr.

### Tissue embedding and sectioning

Individual tissue cassettes were poured with little amount of paraffin at the centre of mold and tissue is placed on it. While the paraffin becomes semi solid the entire cassette is emerged with fresh paraffin and left promptly on top of cold plate to be solidified which is further stored at room temperature in a dry container. To prepare the tissue sections Thermo Shandon microtome was used. Briefly, the embedded tissue blocks were kept on top of ice to rehydrate. Now the blocks were hold in the microtome instrument and sections of 5μm thickness were produced. The paraffin ribbon produced during this step is placed on warm water bath to separate them individually and remove any wrinkles. Now the sections were lifted from water bath using glass slides and kept it to air dry so that it becomes fixed at the surface of the slide.

### Tissue staining with hematoxylin and eosin

In order to stain the tissues with hematoxylin and eosin (H&E) tissue sections were first deparaffinized. For this, slides were initially heated to remove paraffin around the tissue. Further it was washed with xylene for two times 5 min. To rehydrate the tissues it was then treated with 100%, 95%, 70% and 50% ethanol and distilled respectively for 5min. Upon ethanol treatment, the sections were then stained hematoxylin for 1min and washed with distilled water. Thereafter it was stained with eosin for 30sec. Finally, to rehydrate the stained tissue sections were placed into 95% ethanol for 1 min two times and then in 100% ethanol for 2 min thrice. Thereafter the tissues were treated with xylene for 5min thrice. The stained rehydrated tissue sections were then mounted with refrax to observe under the microscope.

### Immunostaining of tissue sections

Similar to the deparaffinization step prior to H&E staining, heated tissue sections were treated with xylene first for 10min initially. Further it was treated with 100% and 70% ethanol respectively for 5min each and then washed with PBS for 5min. Further the tissue slides were placed into sodium citrate buffer (pH 6.0) for 10 min. Now blocking has been performed with 0.2% gelatin for 15 min at room temperature. The samples were further incubated with either anti-Nramp1 or anti-TfR at a dilution of 1: 50 for 2hrs. Further it was washed with PBS and incubated with Alexa tagged secondary antibodies for another 2hrs. Finally after washing the samples with PBS these were mounted with DAPI containing mounting media to counterstain the nuclei.

### Reactive oxygen species (ROS) measurement

To determine the reactive oxygen species within macrophage cells the 2’, 7’-dichlorodihydrofluorescein (H2-DCF) staining technique was used (Shehat & Aranjuez-Tigno, n.d.). Briefly, *L. major* infected or uninfected macrophage cells in presence or absence of ferric ammonium citrate treatment was washed with ice cold PBS properly and fresh medium was added to it containing DCFDA dye at a final concentration of 1μM (dissolved in DMSO). Further the cells were incubated at 37°C at dark for 30min with 5% CO2. Following incubation with the dye media was discarded and cells were washed with ice cold PBS for several times to remove the presence of residual dye. Now the cells are scrapped out of the plate and taken for FACS analysis. Throughout the experiments the cells were kept on ice and in dark. However, in contrast to cellular ROS measurement, the similar parameter within footpad tissues were performed in a different way. Following harvesting the desired tissue from either uninfected or infected mouse, it was incubated with collagenase solution prepared in PBS at a final concentration of 1 mg/ mL at 37° C for half an hour. Thereafter the untangled tissues were taken out of the water bath and a homogenous suspension was prepared using pastel. Further, the tissue suspension was centrifuged at 2000 rpm for 3 min at 4 °C to remove any larger particles present in it. Finally, the supernatant containing homogenous cell suspension is taken and further incubated with DCFDA similarly as mentioned earlier in this section and followed the downstream steps.

### Statistical analysis

All statistical analyses were executed by Student’s t test or one-way ANOVA calculator. The results were represented as mean ± SD from minimum 3 independent experiments. P values of ≤0.05 were considered statistically significant and levels of statistical significance were indicated as follows: * p≤0.05, ** p<0.01, *** p<0.001, **** p<0.0001.

## Acknowledgements

This work was supported by IISER Kolkata intramural fund. S. Banerjee was supported by DST INSPIRE PhD fellowship. The authors sincerely thank Mr. Jajati Keshari Ray and Mr. Sujoy Bose for their expert technical assistance. Drs. Sankar Maiti and Prof. Jayasri Das Sarma are acknowledged for helpful suggestions and for providing critical reagents used in this work.

## Conflict of interests

No competing interests declared.

## Abbreviations

Nramp1: Natural resistance associated macrophage protein 1;
Slc11a1: Solute carrier family 11 member 1;
*L. major*: *Leishmania major*

---

**Figure S1.**
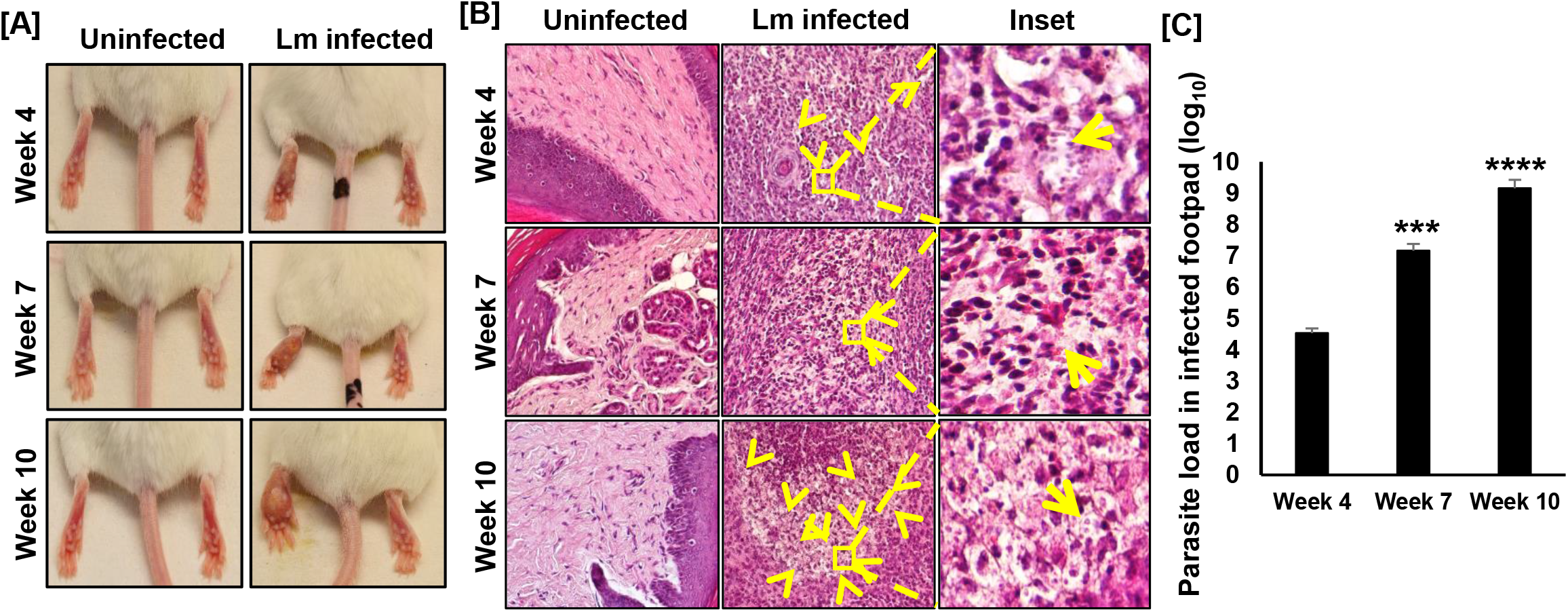
*L. major* infection promotes severe inflammation with the progression of disease. 6-8 weeks old female Balb/c mice were anaesthetized and 1X10^6^ stationary phase metacyclic *L. major* promastigotes were injected into left hind footpad and lesion development was observed upto 10 weeks post infection (p.i.). (A) Representative images of both uninfected and infected mice footpad showing starting from week 4 p.i. the lesion size started to develop significantly till the end of this time course study. (B) At week 4, 7 and 10 post infection mice were euthanized and footpad tissue sections were prepared and further stained with H&E to visualize the tissue architecture of both uninfected and *L. major* (Lm) infected footpad. In the Lm infected panel arrows indicative of the presence of amastigotes with the inset showing enlarged images. Scale bar 10μm. (C) Parasite load in the *L. major* infected mice footpad was determined at week 4, 7 and 10 post infection by limiting dilution assay which is represented in the bar diagram. Error bars represent SEM values calculated from three independent experiments with each experimental group having 5 animals. n.s., non-significant; *p≤ 0.05; **p ≤ .01; ***p≤ 0.001; ****p ≤ .0001.

**Figure S2.**
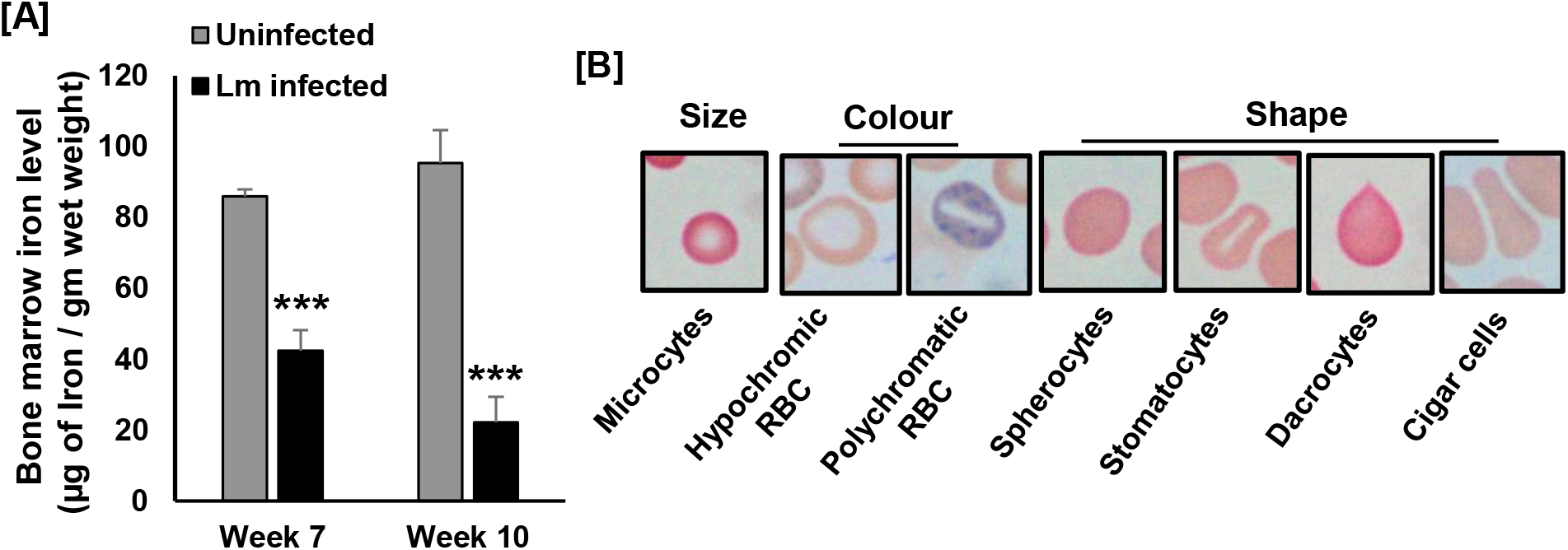
*L. major* infection causes reduced bone marrow iron level with production of damaged RBC. (A) At week 4, 7 and 10 post infection with *L. major* parasites, mice were euthanized from both uninfected and Lm infected groups and bone marrow were harvested from femurs. Further it was lysed properly and lysates from both the mice groups were used to measure the iron content using ferrozine assay and represented in the bar diagram. (B) Blood smear was prepared from the infected mice and stained with Giemsa and further visualized under the microscope. Representative images of different RBCs observed in the Lm infected mice blood smear at week 7 and 10 post infection has been demonstrated.

**Figure S3.**
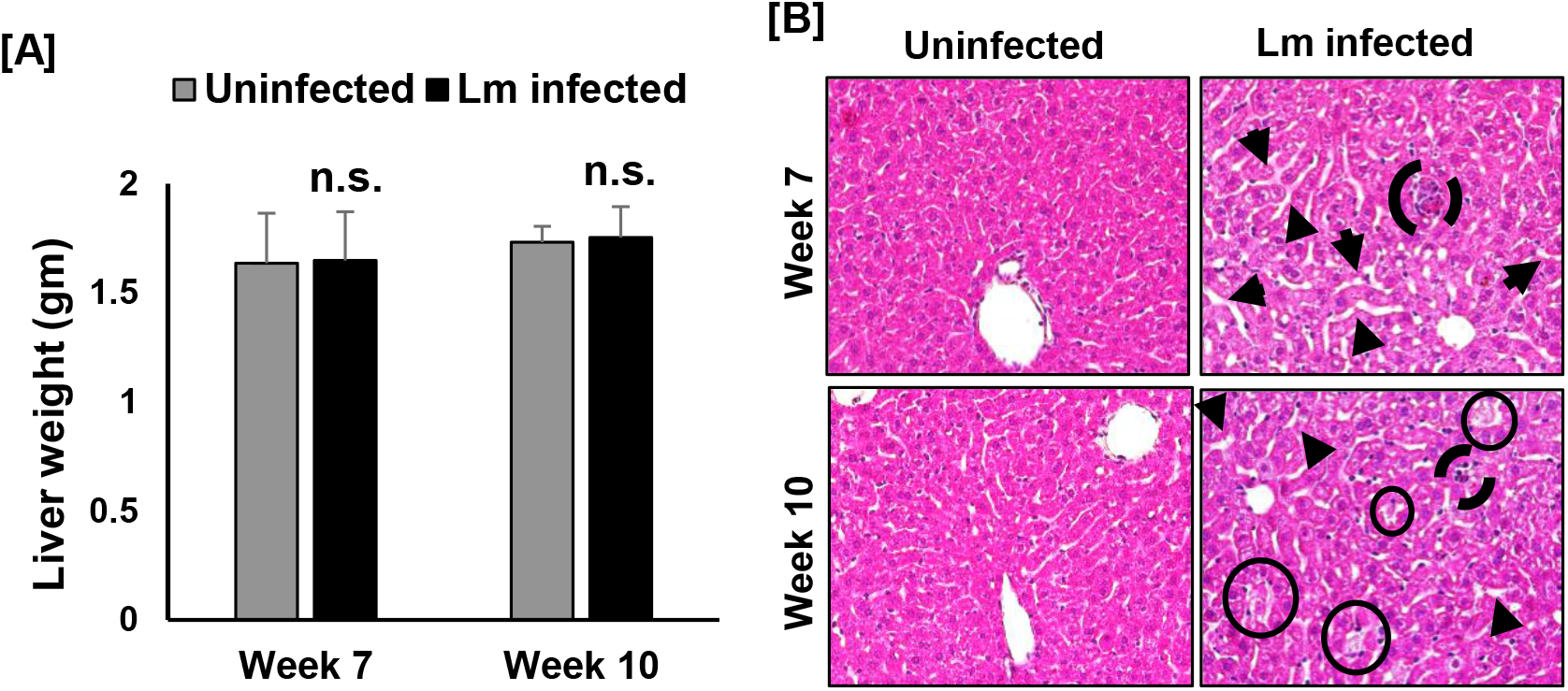
*L. major* infection causes altered hepatic morphology. At week 7 and 10 post infection mice were euthanized and liver was harvested to check any alteration. (A) Liver from different mouse were used to measure the weight and compared in between the uninfected and Lm infected groups which is represented in the bar diagram. Error bars represent SEM values calculated from three independent experiments with each experimental group having at least 5 29 animals. n.s., non-significant; *p≤ 0.05; **p ≤ .01; ***p≤ 0.001; ****p ≤ .0001. (B) Liver tissue sections were prepared and further stained with H&E to visualize the tissue architecture of both uninfected and *L. major* (Lm) infected liver. In the Lm infected panel arrows indicative of the sinusoidal dilation and infiltration of immune cells. Circles indicative of tissue necrosis with the presence of severe inflammation (bold circles). Scale bar 10μm.

**Figure S4.**
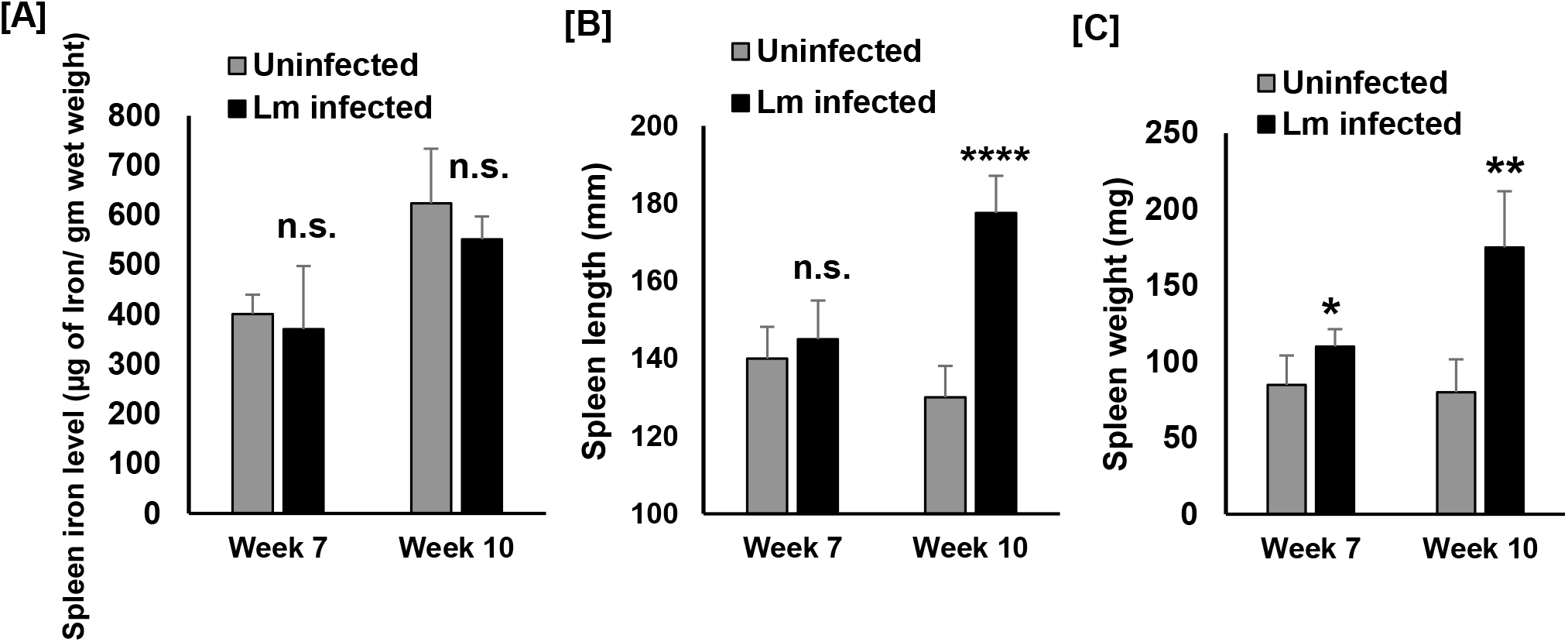
Splenomegaly and iron deficiency are associated with the progression of cutaneous lesion. At week 7 and 10 post infection mice were euthanized and spleen harvested. (A) Spleen lysates were prepared from both uninfected and Lm infected mice groups. Iron content in it was measured using ferrozine assay. Quantification of it is shown the bar diagram. The length (B) as well as weight (C) of both uninfected and Lm infected mice spleen were measured. Bar diagrams represent comparative analysis of these parameters between uninfected (grey bar) and Lm infected (black bar) mice. Error bars represent SEM values calculated from three independent experiments with each experimental group having at least 5 animals. n.s., non-significant; *p≤ 0.05; **p ≤ .01; ****p ≤ .0001.

**Figure S5.**
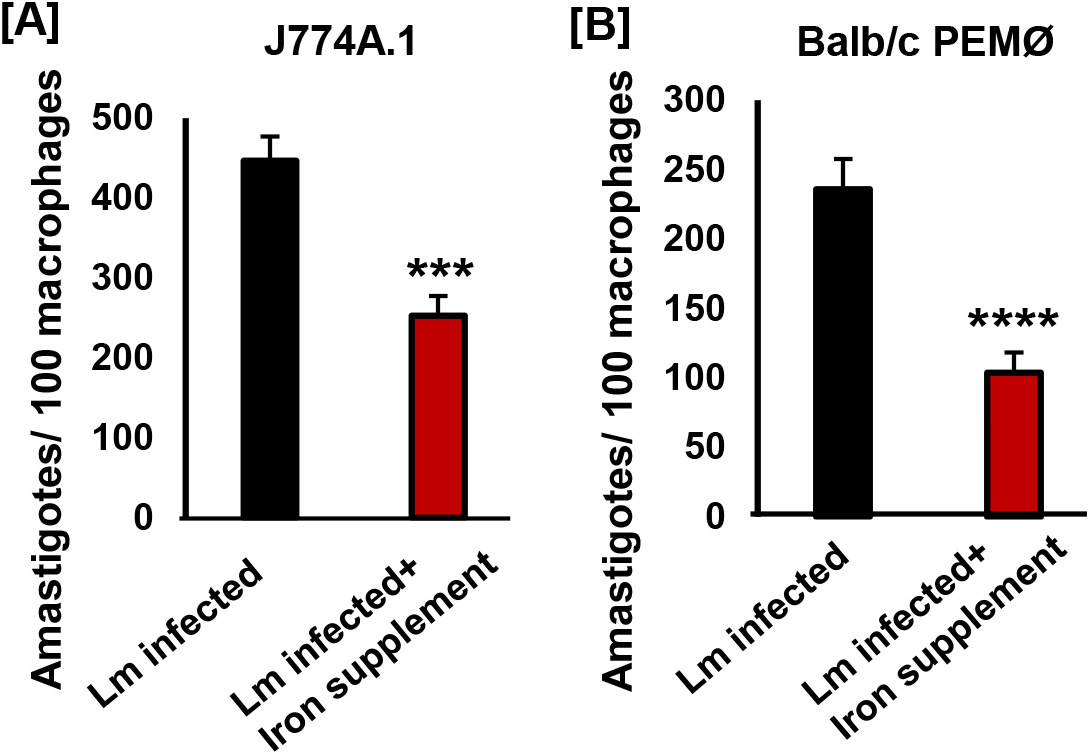
Iron supplementation inhibits *L. major* infection in macrophage cells. Either J774A.1 (A) or Balb/c mouse isolated thioglycollate elicited peritoneal macrophages (B) were infected with *L. major* promastigotes or infected and supplemented with 100μM ferric ammonium citrate. 12hrs post infection, cells were fixed using acetone: methanol and nuclei stained with DAPI to measure the parasite burden in these cells. At least 100 cells from three independent experiments were used to measure the parasite burden and represented in the bar diagram as amastigotes/ 100 macrophage cells. Error bars represent SEM values calculated from three independent experiments. n.s., non-significant; *p≤ 0.05; **p ≤ .01; ***p≤ 0.001; ****p ≤ .0001.

**Figure S6.**
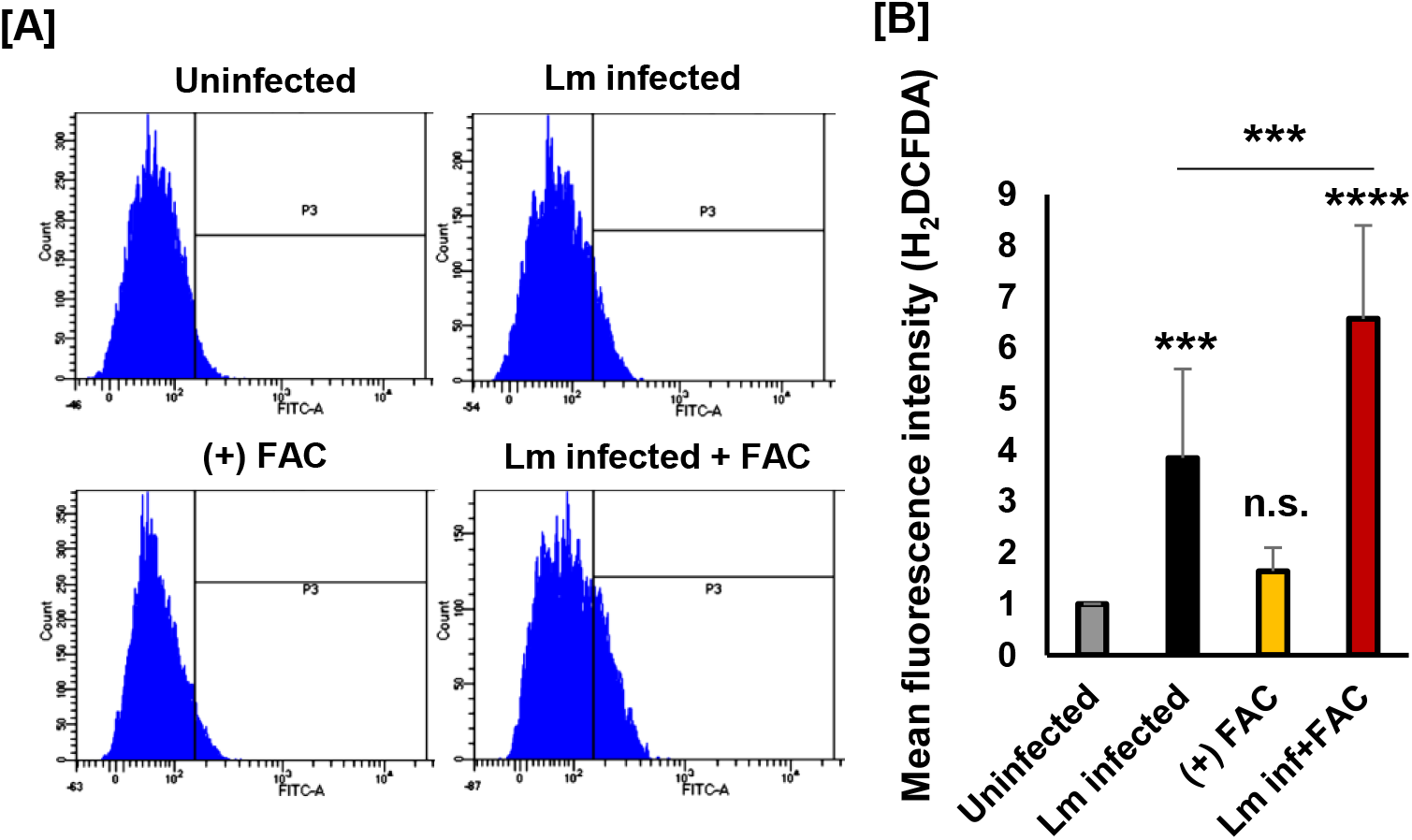
Iron supplementation promotes surge in ROS level in *L. major* infected primary macrophage cells. Balb/c mouse isolated thioglycollate elicited peritoneal macrophages were either uninfected, infected with *L. major* promastigotes, infected and supplemented with 100μM ferric ammonium citrate or supplemented with 100μM ferric ammonium citrate. 12hrs post infection, ROS level was measured using DCFDA based FACS assay (A). (B) Bar diagram is sowing the quantitative analysis of ROS level in those mice groups. Error bars represent SEM values calculated from three independent experiments. ***p≤ 0.001; ****p ≤ .0001.

**Figure S7.**
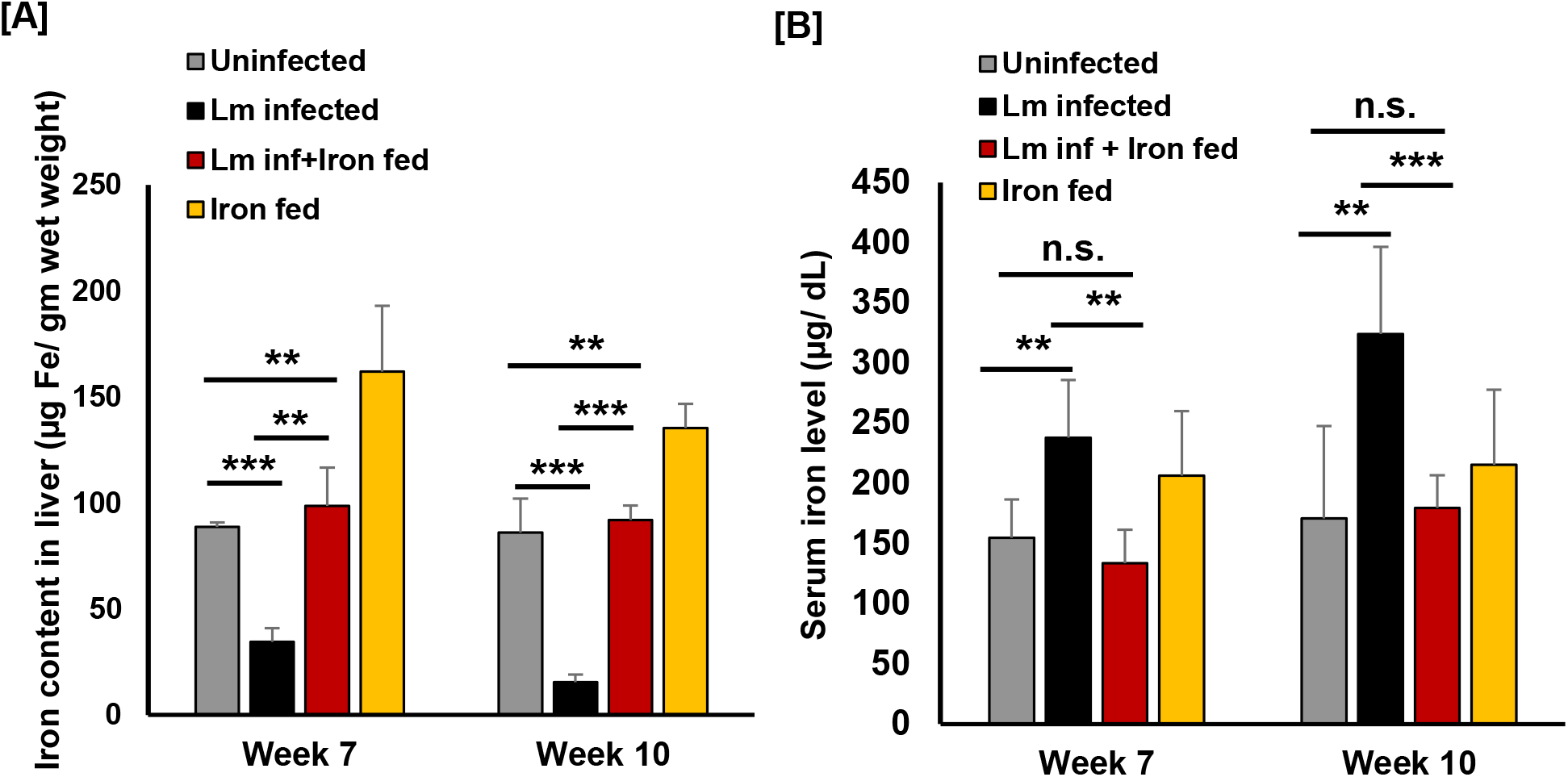
Iron supplementation rescues hepatic iron deficiency and heightened serum iron level in *L. major* infected mouse. From 6-8 weeks old female uninfected, Lm infected, FAC supplemented and Lm infected and FAC supplemented Balb/c mice either blood was collected at week 7 and 10 post infection or euthanized and harvested the livers. Further the livers were used to prepare the lysates and iron content in it was measured using ferrozine assay which is represented in the bar diagram (A). (B) Blood samples were used to prepare the serum and iron level in it was measured using the tibc kit as mentioned in the materials and methods section and represented in the bar diagram. Error bars represent SEM values calculated from three independent experiments with each experimental group having at least 5 animals. n.s., non-significant; *p≤ 0.05; **p ≤ .01; ***p≤ 0.001; ****p ≤ .0001.

## Notes

### Competing Interest Statement

The authors have declared no competing interest.

## References

Abadías-Granado, I., Diago, A., Cerro, P. A., Palma-Ruiz, A. M., & Gilaberte, Y. (2021). Cutaneous and Mucocutaneous Leishmaniasis. Actas Dermo-Sifiliograficas, 112(7), 601–618. https://doi.org/10.1016/j.ad.2021.02.008

Anderson, E. R., & Shah, Y. M. (2013). Iron Homeostasis in the Liver. Comprehensive Physiology, 3(1), 315–330. https://doi.org/10.1002/CPHY.C120016

Banerjee, S., & Datta, R. (2020). Leishmania infection triggers hepcidin-mediated proteasomal degradation of Nramp1 to increase phagolysosomal iron availability. Cellular Microbiology, 22(12), 1–18. https://doi.org/10.1111/cmi.13253

Ben-Othman, R., Flannery, A. R., Miguel, D. C., Ward, D. M., Kaplan, J., & Andrews, N. W. (2014). Leishmania-Mediated Inhibition of Iron Export Promotes Parasite Replication in Macrophages. PLoS Pathogens, 10(1). https://doi.org/10.1371/journal.ppat.1003901

Beutler, E. (2007). Iron storage disease: Facts, fiction and progress. Blood Cells, Molecules, and Diseases, 39(2), 140–147. https://doi.org/10.1016/J.BCMD.2007.03.009

Bisti, S., Konidou, G., Papageorgiou, F., Milon, G., Boelaert, J. R., & Soteriadou, K. (2000). The outcome of Leishmania major experimental infection in BALB/c mice can be modulated by exogenously delivered iron. European Journal of Immunology, 30(12), 3732–3740. https://doi.org/10.1002/1521-4141(200012)30:12<3732::AID-IMMU3732>3.0.CO;2-D

Bose, M., Saha, R., Sen Santara, S., Mukherjee, S., Roy, J., & Adak, S. (2012). Protection against peroxynitrite by pseudoperoxidase from Leishmania major. Free Radical Biology and Medicine, 53(10), 1819–1828. https://doi.org/10.1016/j.freeradbiomed.2012.08.583

Burza, S., Croft, S. L., & Boelaert, M. (2018). Leishmaniasis. The Lancet, 392(10151), 951–970. https://doi.org/10.1016/S0140-6736(18)31204-2

Cairo, G., Bernuzzi, F., & Recalcati, S. (2006). A precious metal: Iron, an essential nutrient for all cells. Genes & Nutrition, 1(1), 25–39. https://doi.org/10.1007/bf02829934

Camaschella, C. (2017). New insights into iron deficiency and iron deficiency anemia. Blood Reviews, 31(4), 225–233. https://doi.org/10.1016/J.BLRE.2017.02.004

Das, N. K., Biswas, S., Solanki, S., & Mukhopadhyay, C. K. (2009). Leishmania donovani depletes labile iron pool to exploit iron uptake capacity of macrophage for its intracellular growth. Cellular Microbiology, 11(1), 83–94. https://doi.org/10.1111/j.1462-5822.2008.01241.x

Das Sarma, J., Fu, L., Tsai, J. C., Weiss, S. R., & Lavi, E. (2000). Demyelination Determinants Map to the Spike Glycoprotein Gene of Coronavirus Mouse Hepatitis Virus. Journal of Virology, 74(19), 9206–9213. https://doi.org/10.1128/jvi.74.19.9206-9213.2000

Donovan, A., Lima, C. A., Pinkus, J. L., Pinkus, G. S., Zon, L. I., Robine, S., & Andrews, N. C. (2005). The iron exporter ferroportin/Slc40a1 is essential for iron homeostasis. Cell Metabolism, 1(3), 191–200. https://doi.org/10.1016/j.cmet.2005.01.003

Dostálová, A., & Volf, P. (2012). Leishmania development in sand flies: Parasite-vector interactions overview. Parasites and Vectors, 5(1), 1–12. https://doi.org/10.1186/1756-3305-5-276

Dumitriu, B., Bhattaram, P., Dy, P., Huang, Y., Quayum, N., Jensen, J., & Lefebvre, V. (2010). Sox6 Is Necessary for Efficient Erythropoiesis in Adult Mice under Physiological and Anemia-Induced Stress Conditions. PLOS ONE, 5(8), e12088. https://doi.org/10.1371/JOURNAL.PONE.0012088

Eruslanov, E. K. (2010). Identification of ROS using DCFDA and flow-cytometry. Oxidative Stress II, 24(1), 6. http://search.ebscohost.com/login.aspx?direct=true%7B&%7Ddb=a9h%7B&%7DAN=36293019%7B&%7Dsite=ehost-live

Ford, J., & Ford, J. (2013). Red blood cell morphology. International Journal of Laboratory Hematology, 35(3), 351–357. https://doi.org/10.1111/IJLH.12082

Ganz, T. (2012). Macrophages and Systemic Iron Homeostasis. Journal of Innate Immunity, 4(5–6), 446–453. https://doi.org/10.1159/000336423

Gurung, P., Karki, R., Vogel, P., Watanabe, M., Bix, M., Lamkanfi, M., & Kanneganti, T. D. (2015). An NLRP3 inflammasome–triggered Th2-biased adaptive immune response promotes leishmaniasis. The Journal of Clinical Investigation, 125(3), 1329–1338. https://doi.org/10.1172/JCI79526

Haas, A. (2007). The phagosome: Compartment with a license to kill. Traffic, 8(4), 311–330. https://doi.org/10.1111/j.1600-0854.2006.00531.x

Huang, H., & Cantor, A. B. (2009). Common features of megakaryocytes and hematopoietic stem cells: What’s the connection? Journal of Cellular Biochemistry, 107(5), 857–864. https://doi.org/10.1002/JCB.22184

Kaye, P., & Scott, P. (2011). Leishmaniasis: Complexity at the host-pathogen interface. Nature Reviews Microbiology, 9(8), 604–615. https://doi.org/10.1038/nrmicro2608

Kulda, J., Poislová, M., Suchan, P., & Tachezy, J. (1999). Iron enhancement of experimental infection of mice by Tritrichomonas foetus. Parasitology Research 1999 85:8, 85(8), 692–699. https://doi.org/10.1007/S004360050617

Lafuse, W. P., Story, R., Mahylis, J., Gupta, G., Varikuti, S., Steinkamp, H., Oghumu, S., & Satoskar, A. R. (2013). Leishmania donovani Infection Induces Anemia in Hamsters by Differentially Altering Erythropoiesis in Bone Marrow and Spleen. PLoS ONE, 8(3), 1–12. https://doi.org/10.1371/journal.pone.0059509

Liao, C., Hardison, R. C., Kennett, M. J., Carlson, B. A., Paulson, R. F., & Prabhu, K. S. (2018). Selenoproteins regulate stress erythroid progenitors and spleen microenvironment during stress erythropoiesis. Blood, 131(23), 2568–2580. https://doi.org/10.1182/BLOOD-2017-08-800607

Luck, A. N., & Mason, A. B. (2012). Transferrin-Mediated Cellular Iron Delivery. Current Topics in Membranes, 69, 3–35. https://doi.org/10.1016/B978-0-12-394390-3.00001-X

Mackenzie, E. L., Iwasaki, K., & Tsuji, Y. (2008). Comprehensive Invited Review Intracellular Iron Transport and Storage: From Molecular Mechanisms to Health Implications. ANTIOXIDANTS & REDOX SIGNALING, 10(6). https://doi.org/10.1089/ars.2007.1893

Maltezou, H. C. (2010). Drug resistance in visceral leishmaniasis. Journal of Biomedicine and Biotechnology, 2010. https://doi.org/10.1155/2010/617521

Melo, C. V. B. de, Hermida, M. D. E. R., Mesquita, B. R., Fontes, J. L. M., Koning, J. J., Solcà, M. da S., Benevides, B. B., Mota, G. B. S., Freitas, L. A. R., Mebius, R. E., & dos-Santos, W. L. C. (2020). Phenotypical Characterization of Spleen Remodeling in Murine Experimental Visceral Leishmaniasis. Frontiers in Immunology, 11, 653. https://doi.org/10.3389/FIMMU.2020.00653/BIBTEX

Nairz, M., Haschka, D., Demetz, E., & Weiss, G. (2014). Iron at the interface of immunity and infection. Frontiers in Pharmacology, 5 JUL, 152. https://doi.org/10.3389/FPHAR.2014.00152/BIBTEX

Navas, S., Seo, S.-H., Chua, M. M., Sarma, J. Das, Lavi, E., Hingley, S. T., & Weiss, S. R. (2001). Murine Coronavirus Spike Protein Determines the Ability of the Virus To Replicate in the Liver and Cause Hepatitis. Journal of Virology, 75(5), 2452–2457. https://doi.org/10.1128/jvi.75.5.2452-2457.2001

Nemeth, E., & Ganz, T. (2006). Regulation of Iron Metabolism by Hepcidin. http://Dx.Doi.Org/10.1146/Annurev.Nutr.26.061505.111303, 26, 323–342. https://doi.org/10.1146/ANNUREV.NUTR.26.061505.111303

Palumbo, E. (2010). Treatment strategies for mucocutaneous leishmaniasis. Journal of Global Infectious Diseases, 2(2), 147. https://doi.org/10.4103/0974-777x.62879

Riemer, J., Hoepken, H. H., Czerwinska, H., Robinson, S. R., & Dringen, R. (2004). Colorimetric ferrozine-based assay for the quantitation of iron in cultured cells. Analytical Biochemistry, 331(2), 370–375. https://doi.org/10.1016/j.ab.2004.03.049

Sacks, D. L., & Melby, P. C. (2015). Animal models for the analysis of immune responses to leishmaniasis. In Current Protocols in Immunology (Vol. 2015). https://doi.org/10.1002/0471142735.im1902s108

Shehat, M. G., & Aranjuez-Tigno, J. (n.d.). Flow Cytometric Measurement Of ROS Production In Macrophages In Response To FcγR Cross-linking. https://www.jove.com/video/59167/

Siatecka, M., & Bieker, J. J. (2011). The multifunctional role of EKLF/KLF1 during erythropoiesis. Blood, 118(8), 2044–2054. https://doi.org/10.1182/BLOOD-2011-03-331371

Soares, M. P., & Hamza, I. (2016). Macrophages and Iron Metabolism. Immunity, 44(3), 492–504. https://doi.org/10.1016/J.IMMUNI.2016.02.016

Soe-Lin, S., Apte, S. S., Andriopoulos, B., Andrews, M. C., Schranzhofer, M., Kahawita, T., Garcia-Santos, D., & Ponka, P. (2009). Nramp1 promotes efficient macrophage recycling of iron following erythrophagocytosis in vivo. Proceedings of the National Academy of Sciences of the United States of America, 106(14), 5960–5965. https://doi.org/10.1073/pnas.0900808106

Steverding, D. (2017). The history of leishmaniasis. Parasites and Vectors, 10(1), 1–10. https://doi.org/10.1186/s13071-017-2028-5

Taylor, M. C., & Kelly, J. M. (2010). Iron metabolism in trypanosomatids, and its crucial role in infection. Parasitology, 137(6), 899–917. https://doi.org/10.1017/S0031182009991880

Theurl, I., Hilgendorf, I., Nairz, M., Tymoszuk, P., Haschka, D., Asshoff, M., He, S., Gerhardt, L. M. S., Holderried, T. A. W., Seifert, M., Sopper, S., Fenn, A. M., Anzai, A., Rattik, S., McAlpine, C., Theurl, M., Wieghofer, P., Iwamoto, Y., Weber, G. F., … Swirski, F. K. (2016). On-demand erythrocyte disposal and iron recycling requires transient macrophages in the liver. Nature Medicine 2016 22:8, 22(8), 945–951. https://doi.org/10.1038/nm.4146

Upton, R. L., Chen, Y., Mumby, S., Gutteridge, J. M. C., Anning, P. B., Nicholson, A. G., Evans, T. W., & Quinlan, G. J. (2003). Variable tissue expression of transferrin receptors: relevance to acute respiratory distress syndrome. European Respiratory Journal, 22(2), 335–341. https://doi.org/10.1183/09031936.03.00075302

Wang, H., Liu, D., Song, P., Jiang, F., Chi, X., & Zhang, T. (2021). Exposure to hypoxia causes stress erythropoiesis and downregulates immune response genes in spleen of mice. BMC Genomics, 22(1), 1–14. https://doi.org/10.1186/S12864-021-07731-X/FIGURES/5

Wang, S., He, X., Wu, Q., Jiang, L., Chen, L., Yu, Y., Zhang, P., Huang, X., Wang, J., Ju, Z., Min, J., & Wang, F. (2020). Transferrin receptor 1-mediated iron uptake plays an essential role in hematopoiesis. Haematologica, 105(8), 2071–2082. https://doi.org/10.3324/HAEMATOL.2019.224899

Weigel, M. M., Armijos, R. X., Zurita, C., Racines, J., Reddy, A., & Mosquera, J. (1995). Nutritional status and cutaneous leishmaniasis in rural Ecuadorian children. Journal of Tropical Pediatrics, 41(1), 22–28. https://doi.org/10.1093/tropej/41.1.22

Zabolotzky, S. M., & Walker, D. B. (1990). Peripheral Blood Smear. Cowell and Tyler’s Diagnostic Cytology and Hematology of the Dog and Cat, 438–467. https://doi.org/10.1016/b978-0-323-53314-0.00026-2

Zhou, Y., Que, K. T., Zhang, Z., Yi, Z. J., Zhao, P. X., You, Y., Gong, J. P., & Liu, Z. J. (2018). Iron overloaded polarizes macrophage to proinflammation phenotype through ROS/acetyl-p53 pathway. Cancer Medicine, 7(8), 4012–4022. https://doi.org/10.1002/CAM4.1670

Zulfiqar, B., Shelper, T. B., & Avery, V. M. (2017). Leishmaniasis drug discovery: recent progress and challenges in assay development. Drug Discovery Today, 22(10), 1516–1531. https://doi.org/10.1016/j.drudis.2017.06.004

